# RNA G-Quadruplexes Function as a Tunable Switch of FUS Phase Separation

**DOI:** 10.1101/2025.10.31.685846

**Authors:** Jenny L. Carey, Miyuki Hayashi, Emily Welebob, Laura R. Ganser, Huan Wang, Kerry Buckhaults, Jacquelyn A. DePierro, Zheng Shi, James Shorter, Sua Myong, Aaron R. Haeusler, Lin Guo

## Abstract

FUS undergoes liquid-liquid phase separation (LLPS) to support essential cellular functions, but aberrant phase transitions promote toxic aggregation in neurodegenerative disease. Short RNA oligonucleotides can reverse this behavior, yet the structural determinants that govern RNA activity remain poorly defined. Here, we identify RNA G-quadruplexes (rG4s) as tunable structural motifs that potently modulate FUS LLPS. rG4 activity depends on its concentration and is modulated by rG4 length and stability: increasing repeat number switches rG4s from inhibitor to nucleator of FUS assembly, whereas chemical modifications that stabilize rG4 enhance inhibitory function and render these activities resilient to ionic perturbation. Although short rG4s interact with both soluble and condensed FUS, they preferentially engage the soluble pool, likely shifting the equilibrium toward dispersion. Leveraging these mechanistic insights, we developed a bioinformatic pipeline that uncovered more rG4 inhibitors that robustly reverse FUS LLPS and aggregation. Our findings establish rG4s as chemically programmable regulators of protein phase behavior and provide a blueprint for engineering RNA-based therapeutics that dissolve pathogenic FUS assemblies. More broadly, this work directly links RNA secondary structure to distinct functional outcomes in phase behavior, establishing a structure-function paradigm for RNA control of condensates, demonstrating implications in both fundamental biology and therapeutic development.

**GRAPHICAL ABSTRACT:** 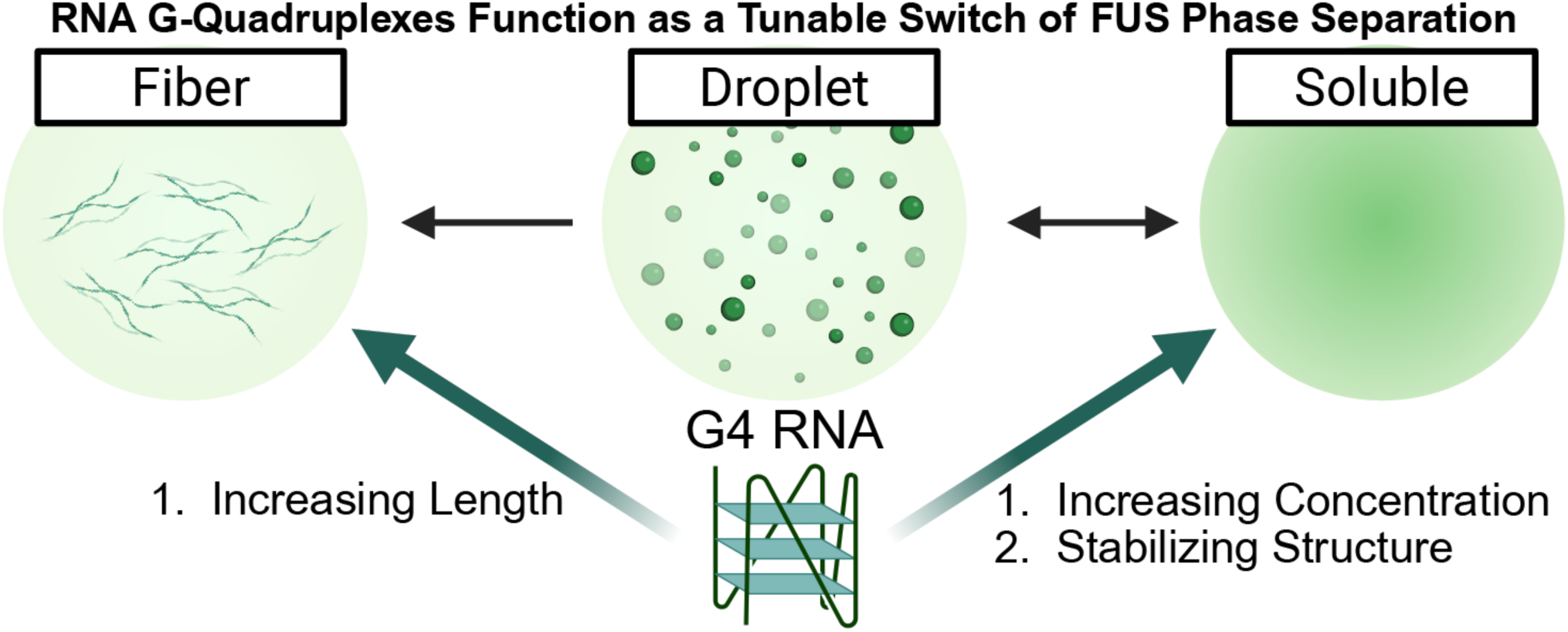

## INTRODUCTION

Liquid-liquid phase separation (LLPS) of RNA-binding proteins (RBPs) and RNAs has emerged as a fundamental mechanism for organizing cellular components into dynamic biomolecular condensates supporting a wide range of biological processes, yet the mechanisms that regulate RBP phase behavior are not fully understood. Among these RBPs, Fused in Sarcoma (FUS) is one of the most extensively studied and serves as a key model system for understanding how RNA and protein features control LLPS. FUS plays essential roles in key cellular processes, including transcriptional regulation, RNA splicing, and genomic stability maintenance^1,2^. A hallmark feature of FUS is its ability to undergo LLPS, primarily driven by its N-terminal prion-like domain (PrLD) and C-terminal RGG domains^3–7^. More recent studies have shown that additional structured elements, including RNA-binding domains, also regulate FUS LLPS^8,9^. Phase separation enables FUS to assemble into membraneless organelles, such as stress granules (SGs), and is essential for its physiological function^10^. However, dysregulation of FUS LLPS can trigger a pathological phase transition, where liquid-like condensates convert into solid-like aggregates and fibrils^4,11–13^. This aberrant behavior is a hallmark of neurodegenerative diseases, including amyotrophic lateral sclerosis (ALS) and frontotemporal dementia (FTD), and is exacerbated by disease-linked mutations in FUS^4,13,14^. Understanding the molecular mechanisms that regulate FUS phase behavior is therefore crucial for developing strategies to block or reverse toxic protein aggregation.

Multiple cellular factors regulate FUS phase separation. Nuclear import receptors (NIRs), for example, modulate LLPS by binding the proline-tyrosine nuclear localization signal (PY-NLS) of FUS, offering a promising therapeutic mechanism of action^15–19^. Post-translational modifications, such as phosphorylation and acetylation, can also modulate FUS LLPS^12,17,20,21^. In addition, physicochemical features of the cellular environment, such as ionic strength^9,22^, temperature^23,24^, and crowding^25^, can influence FUS phase behavior. Various FUS-binding biomolecules further contribute to this regulation. For instance, poly(ADP-ribose) (PAR) promotes FUS LLPS^26^. RNA likewise modulates FUS condensation and displays a biphasic effect on FUS LLPS^27–29^. It was reported that at low concentrations, RNA enhances FUS LLPS^3^, whereas at high concentrations, such as in the mammalian nucleus, RNA suppresses condensation by maintaining FUS in a soluble state^27^.

The broad sequence and structural diversity of RNA, along with its ability to either promote or inhibit FUS LLPS^27–29^, makes RNA an attractive platform for therapeutic development. The large sequence space allows for the identification of specific RNA oligonucleotides that can effectively modulate FUS LLPS and aberrant phase transitions. In previous work, we identified short RNAs with distinct functional profiles, including strong inhibitors that fully suppress both FUS LLPS and aggregation, and weak inhibitors that prevent aggregation but preserve liquid droplets^30^. Notably, the strong inhibitor, RNA S1, mitigates aberrant FUS phase behavior in neurons and enhances the survival of disease neurons harboring ALS-causing FUS mutation^30^, underscoring therapeutic potential. However, the mechanistic basis for these differential RNA activities remains poorly understood. In particular, it is unclear why only a subset of RNAs are effective inhibitors. This knowledge gap limits rational design of RNA-based therapeutics and constrains our understanding of how RNAs regulate FUS LLPS in cells.

In this study, we uncover the structural principles by which short RNAs regulate FUS phase separation, identifying RNA G-quadruplexes (rG4s) as key determinants of inhibitory activity. rG4-containing RNAs suppress FUS LLPS in a concentration-dependent manner, and their function can be precisely tuned by modifying structural features. Increasing rG4 repeat number switches inhibitory RNAs into nucleators of FUS assembly, while chemical stabilization of rG4 enhances potency and confers strong activities that are resistant to ionic stress. Guided by these principles, we developed a bioinformatic pipeline to search the human transcriptome and identified a new class of endogenous rG4s that robustly reverse FUS LLPS and aggregation. Together, these findings establish rG4 as a programmable structural code for modulating protein condensation and provide a design framework for engineering structured RNAs with therapeutic potential. More broadly, this work advances our fundamental understanding of the mechanistic rules by which RNA structure governs protein condensate dynamics, establishing a structure-function paradigm for RNA-mediated phase regulation with broad biological and therapeutic implications.

### MATERIALS AND METHODS

## RNA Production

The S1*10 expression vector, pUC-GW-Kan-rpRNA2, was synthesized by GENEWIZ from Azenta Life Sciences. To generate S1*10, plasmid was digested with EcoRV-HF (NEB, R3195S) and SnaBI (NEB, R0130S) for 16 h at 37°C. Digested product was run on a 1% agarose gel and the ∼250 base-pair fragment was isolated using QIAquick Gel Extraction Kit (QIAGEN, 28706) according to manufacturer protocol. The DNA fragment was concentrated using QIAquick PCR Purification Kit (QIAGEN, 28106) and eluted with Ultra Pure water instead of kit-provided Elution Buffer as the presence of EDTA could prevent T7 RNA polymerase activity. In vitro transcription was performed using TranscriptAid T7 High Yield Transcription Kit (Thermo Scientific, K0441) at 37°C for 16 h. RNA S1*10 was purified from the reaction using acidic phenol-chloroform extraction followed by RNA Clean & Concentrator-100 Kit (Zymo, R1019), following the “Purification of Small and Large RNAs into Separate Fractions” protocol, provided by the manufacturer. The Large RNA fraction (S1*10) was concentrated using RNA Clean & Concentrator-5 Kit (Zymo, R1013). RNA S1*10 was analyzed by urea-containing denaturing gel and stained with ethidium bromide to verify purity.

All other oligonucleotides were synthesized by Horizon Discovery. For tagged RNAs, fluorescein-, Cy5-, or Cy3-dyes were conjugated to the 5’ end of the indicated oligonucleotide. The sequences of all oligonucleotides used are listed in **Table S1**. G-Quadruplex prediction was obtained using the first iteration of QGRS Mapper^31^ (https://bioinformatics.ramapo.edu/QGRS/index.php) using the following parameters: QGRS Max Length of 30, Minimum G-Group Size of 2, and Loop Size from 0-36. Canonical RNA structure prediction was obtained using RNAfold server^32^. If the oligonucleotide was predicted to form canonical secondary structure, the predicted minimum free energy (MFE) structure frequency was recorded in **Table S1**.

## Protein Expression and Purification

### MBP-FUS and MBP-FUS-GFP Purification

Plasmids encoding MBP-TEV-FUS^18^ or 6xHis-MBP-TEV-FUS-GFP^14^ were expressed in BL21-CodonPlus (DE3)-RIL cells (Agilent, 230245) and protein expression was induced with 1 mM IPTG for 16 h at 15°C in the presence of dextrose. Cells were pelleted and resuspended in Wash Buffer (20 mM HEPES-NaOH pH 7.5, 50 mM NaCl, 2 mM EDTA, 10% Glycerol, and 2 mM DTT) supplemented with cOmplete EDTA-free Protease Inhibitor Cocktail (Roche, 5056489001). Resuspended cells were lysed with 0.2 mg/mL lysozyme (Gold Bio, L-040-25) followed by sonication. Affinity purification was performed using Amylose Resin (NEB, E8021L) and protein was eluted in MBP-FUS Elution Buffer (20 mM HEPES-NaOH pH 7.5, 50 mM NaCl, 2 mM EDTA, 10% Glycerol, 10 mM maltose, and 2 mM DTT). Eluted protein was further purified by HiTrap Heparin HP affinity column (Cytiva, 17040703) with a linear gradient of Low-Salt Wash Buffer (50 mM HEPES-NaOH pH 7.4, 50 mM NaCl, 2 mM EDTA, 10% Glycerol, and 2 mM DTT) and High-Salt Wash Buffer (50 mM HEPES-NaOH pH 7.4, 1 M NaCl, 2 mM EDTA, 10% Glycerol, and 2 mM DTT) to remove RNA contamination. Protein purity was assessed by SDS-PAGE with Coomassie staining, and pure fractions were pooled for experiments. MBP-FUS was purified to an OD260/280 ratio of ∼0.6 to ensure the removal of any co-purifying RNA.

### GST-FUS Purification

GST-FUS was purified according to previously published methods^15,33^. Briefly, pGST-Duet-FUS^33^ was expressed in BL21-CodonPlus (DE3)-RIL cells (Agilent, 230245) and induced with 1 mM IPTG for 16 h at 15°C. Cells were resuspended in PBS supplemented with cOmplete EDTA-free Protease Inhibitor Cocktail (Roche, 5056489001), lysed with sonication, and purified with Glutathione Sepharose 4 Fast Flow Media (Cytiva, 17513201). Protein was eluted with FUS Assembly Buffer (50 mM Tris-HCl, pH 8.0, 200 mM trehalose, and 20 mM glutathione).

### pHis-TEV Purification

TEV was purified according to previously published methods^34^. Briefly, pHis-TEV was expressed in BL21-CodonPlus (DE3)-RIL cells (Agilent, 230245) and induced with 1 mM IPTG for 16 h at 15°C. Cells were resuspended in Lysis Buffer (25 mM Tris-HCl pH 8.0, 500 mM NaCl, and 10 mM β -ME) supplemented with cOmplete EDTA-free Protease Inhibitor Cocktail (Roche, 5056489001), lysed with sonication, and affinity purified with HisPur Ni-NTA Resin (Thermo, 88223). Protein was eluted with TEV Elution Buffer (25 mM Tris-HCl pH 8.0, 500 mM NaCl, 300 mM Imidazole, and 10 mM β -ME) and dialyzed overnight into Dialysis Buffer (25 mM Tris-HCl pH 8.0, 5% Glycerol, 5 mM β -ME). Dialyzed TEV was further purified by HiTrap SP XL column (Cytiva, 17-5160-1) with a linear gradient of TEV Low Salt Buffer (25 mM HEPES-NaOH pH 7.0, 5% glycerol, and 5 mM DTT) and TEV High Salt Buffer (25 mM HEPES-NaOH pH 7.0, 750 mM NaCl, 5% glycerol, and 5 mM DTT). Purified TEV was stored in TEV High Salt Buffer containing 50% glycerol at 2 mg/mL concentration. Activity of MBP-FUS cleavage by purified TEV was validated before use.

## Phase Separation Assays

MBP-FUS and MBP-FUS-GFP were buffer exchanged into Turbidity Assay Buffer (TAB; 50 mM Tris-HCl pH 7.4, 0.5% Glycerol) supplemented with 100 mM KCl or 100 mM NaCl (buffer system referred to as “TAB-K^+^” or “TAB-Na^+^”) using 10 mL Zeba 40K MWCO Spin Desalting Columns (Thermo Scientific, A57765) according to manufacturer protocol. Prior to all experiments, MBP-FUS and MBP-FUS-GFP were centrifuged for 10 minutes at 21,000 g and 4°C, and the clear supernatant was used to obtain protein concentration using Bradford assay.

Phase separation assays in Figures 1 and S1 were performed in TAB-Na^+^ and did not use folded RNA. For phase separation assays comparing the effects of magnesium (**Figs. S2C-D, F-H, J-K; 5G-H**), magensium was added to MBP-FUS in TAB-K^+^ to a final concentration of 3 mM and did not use folded RNA. For the remaining phase separation assays, MBP-FUS buffer is noted in the figure legends and folded RNA was used. To fold RNA, RNAs were diluted in RNA Folding Buffer (10 mM Tris-HCl pH 7.4 supplemented with 50 mM KCl or 50 mM NaCl) and heated at 90°C for 3 minutes followed by cooling to 25°C at a rate of 1°C/min in a thermal cycler. When multiple RNAs are added to the same well (**Fig. 6A-B**), RNAs were folded independently to prevent intramolecular interactions, then mixed immediately prior to adding to pre-formed FUS droplets.

**Figure 1.**
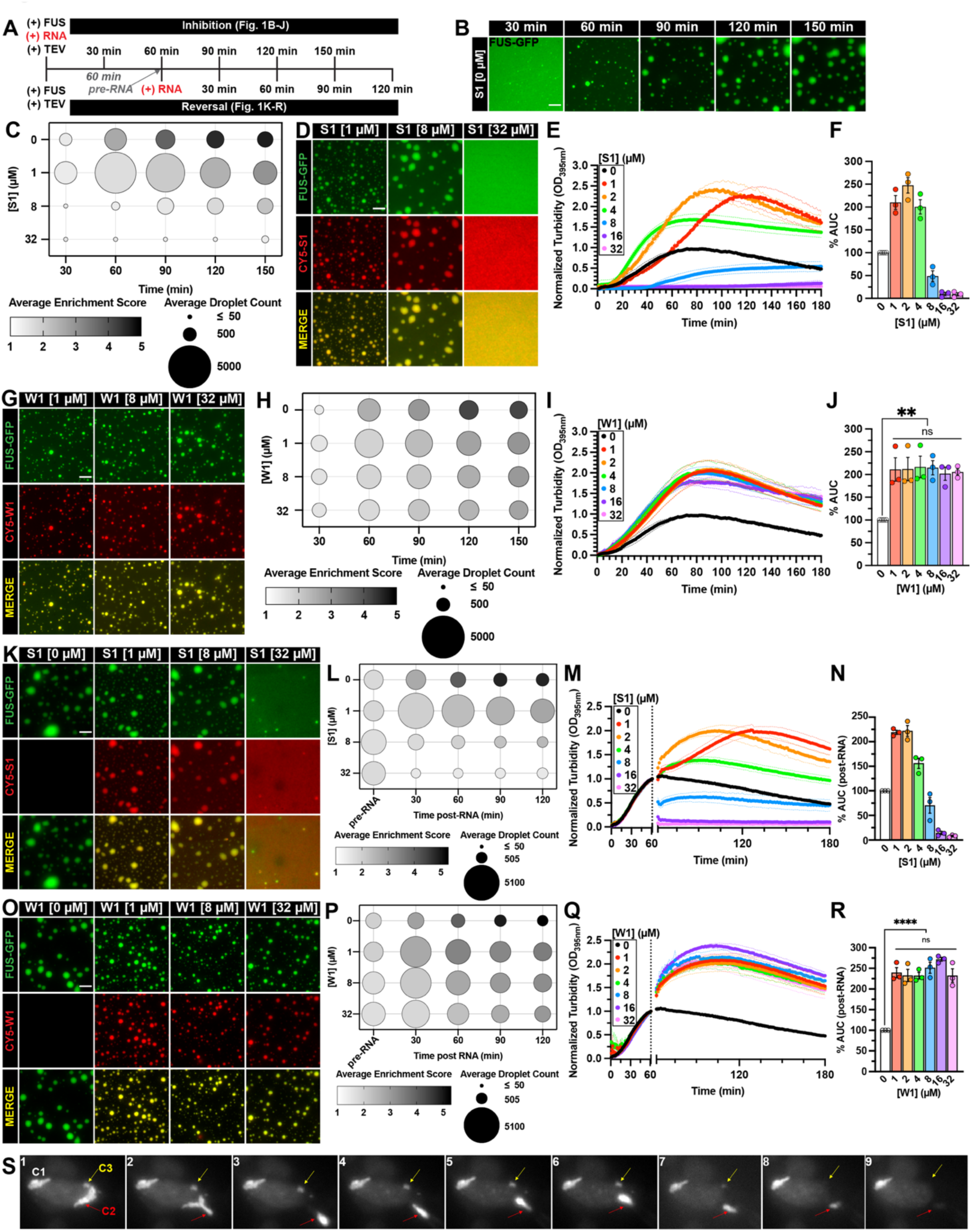
Strong RNA inhibitors prevent and reverse FUS phase separation in a concentration-dependent manner. **(A)** Schematic of experimental paradigms. FUS phase separation is initiated by adding TEV protease to cleave MBP tag from MBP-TEV-FUS. For inhibition experiments, RNA is added to FUS phase separation reactions prior to initiating FUS phase separation with TEV protease. For reversal experiments, FUS phase separation is initiated by TEV protease 60 minutes prior to adding RNA. **(B)** Representative images of FUS droplets formed without RNA. LLPS of FUS (1.8 µM FUS supplemented with 0.2 µM FUS-GFP) was monitored by droplet imaging at indicated times. Scale bar is 5 µm. The images without RNA presented in Fig. 1B were collected in the same trial as the data presented in Fig. 1D. (**C, H**) Phase diagrams summarizing the inhibition of FUS LLPS by RNA S1 (**C**) or W1 (**H**). Average droplet count defines symbol size, and average FUS enrichment score (quantified by dividing mean GFP signal within droplets by mean GFP signal in background) defines symbol saturation. Data used to generate phase diagrams are the mean of three independent experiments. (**D, G**) Representative images of FUS droplets formed in the presence of RNA S1 (**D**) or W1 (**G**). LLPS of FUS (1.8 µM FUS supplemented with 0.2 µM FUS-GFP) with RNA (1, 8, or 32 µM unlabeled RNA supplemented with 75 nM Cy5-RNA) was monitored by droplet imaging every 30 minutes for a duration of 150 minutes. 120-minute timepoint images were shown due to space limitations to represent RNA effect on FUS LLPS. Scale bar is 5 µm. (**E, I**) Turbidity measurement of FUS LLPS inhibition by RNA S1 (**E**) or W1 (**I**). MBP-FUS (2 µM) was incubated in the presence or absence of RNA prior to initiating LLPS reaction with TEV protease. Turbidity at 395 nm was monitored at 25°C over 180 min. Data represent mean±SEM (n=3-4). The 0 µM RNA control plotted in Figs. 1E, 1I, and S1C are the same because these experiments were conducted simultaneously. (**F, J**) Area under the curve (AUC) was calculated from turbidity curves in Fig. 1E (**F**) and Fig. 1I (**J**). Data represent mean±SEM (n=3-4). (**J**) Ordinary one-way ANOVA with Tukey’s multiple comparisons test was used to compare the means between RNA W1 concentrations (ns P>0.05, **P≤0.005). (**K, O**) Representative images of FUS droplets 60 minutes after the addition of RNA S1 (**K**) or W1 (**O**). FUS (1.8 µM FUS supplemented with 0.2 µM FUS-GFP) droplets were formed for 60 minutes prior to adding RNA (1, 8, or 32 µM unlabeled RNA supplemented with 75 nM Cy5-RNA). Droplets were monitored by imaging immediately before RNA, then every 30 minutes for a duration of 120 minutes after RNA was added. The 60-minute post-RNA timepoint images were selected to represent RNA’s effect on FUS LLPS. Scale bar is 5 µm. (**L, P**) Phase diagrams summarizing the reversal of FUS LLPS by RNA S1 (**L**) or W1 (**P**). Droplet images collected in Fig. 1K (**L**) or 1O (**P**) were quantified. Average droplet count defines symbol size and average FUS enrichment score defines symbol saturation. Data used to generate phase diagrams are the mean of three independent experiments. (**M, Q**) Turbidity measurement of FUS LLPS reversal with RNA S1 (**M**) or W1 (**Q**). FUS (2 µM) LLPS was initiated, and monitored by turbidity at 395 nm. After 60 min, data collection was paused to add RNA then resumed to monitor turbidity changes for an additional 120 min. Data represent mean±SEM (n=3). The 0 µM RNA control plotted in Fig. 1M and 1Q are the same because these experiments were conducted simultaneously. (**N, R**) AUC was calculated from turbidity curves in Fig. 1M (**N**) and Fig. 1Q (**R**) after RNA was added. Data represent mean±SEM (n=3). (**R**) Ordinary one-way ANOVA with Tukey’s multiple comparisons test was used to compare the means between RNA W1 concentrations (ns P>0.05, ****P<0.0001). (**S**) Microinjection of RNA S1 (100 µM) into HEK 293T cells expressing FUS-WT-eGFP. Panel 1 shows pre-injection condensates 1 (C1), 2 (C2, red arrows), and 3 (C3, yellow arrows). Panels 2-3 show suction pressure, where C2 flows into the pipette tip. Panels 4-6 show ejection pressure (∼10 kPa), where C2 is blocking RNA injection. Panels 7-8 show C2 and C3 dissolving after RNA S1 injection. Panel 9 is after stopping RNA injection.

### Droplet Imaging

In a glass-bottom 96-well plates (MatTek, PBK96G-1.5-5-F), droplet reactions were prepared in TAB-K^+^ or TAB-Na^+^ (1.8 µM MBP-FUS, 0.2 µM MBP-FUS-GFP, and 10 mM DTT). When RNA was tested, 75 nM Cy5-labeled RNA was included to the indicated concentration of unlabeled RNA to aid in visualization. TEV protease was used at 8 μg/mL to initiate MBP-FUS phase separation. For inhibition assays, RNA was added prior to TEV and droplet formation was monitored. For reversal assays, TEV protease was added to initiate MBP-FUS phase separation. FUS was allowed to form droplets for 60-90 minutes prior to adding RNA or appropriate buffer control. For the S1*10 droplet assay (**Figs. 4J-K, S5B-C**), droplet reactions were prepared in TAB-K^+^ (500 nM MBP-FUS-GFP in TAB-K^+^, indicated RNA, 10 mM DTT, and 8 µg/mL TEV) and sandwiched between two glass coverslips (Epredia, 102455) using SecureSeal Spacers (Electron Microscopy Sciences, 70327-9S).

To monitor droplet formation or reversal, images were acquired using 100x/1.4-0.7 oil objective on a DMi8 microscope (Leica, Germany). For each condition, images were acquired in 30-min intervals at 10 pre-defined regions across the entire well. All image analysis was performed in Fiji^35^. Briefly, a lower-bound threshold was applied to each image to select all droplets after which the size, area, and mean fluorescence intensity were measured. Enrichment score was calculated by dividing the mean fluorescence intensity of the signal inside each droplet by the mean fluorescence intensity of the background.

### Turbidity Assays

Turbidity reactions were prepared in 96-well plates (Thermo Scientific, 265301). Briefly, turbidity reactions were prepared in TAB-K^+^ or TAB-Na^+^ (2 µM MBP-FUS and 10 mM DTT) and TEV protease (8 µg/mL) was added to each reaction to initiate FUS phase separation. Unlabeled RNAs were used to assess their ability to inhibit or reverse FUS phase separation. Turbidity measurements were collected at 395 nm in 1-minute cycles at 25°C using TECAN Spark plate reader. For each trial, the average buffer turbidity was subtracted from all curves. For inhibition assays, all curves were normalized to the maximum absorbance of the 0 µM RNA control. Area under the curve (AUC) was quantified for all curves and normalized to the 0 µM RNA control AUC for each trial. For reversal assays, all curves were normalized to their individual pre-RNA absorbance in order to assess relative response to RNA addition. AUC was quantified after RNA addition for all curves and normalized to the 0 µM RNA AUC in each buffer condition.

## FUS Disaggregation and Sedimentation Analysis

Prior to experiments, GST-FUS was centrifuged for 10 minutes at 21,000 g and 4°C, and the clear supernatant was used to obtain protein concentration using Bradford assay. FUS aggregation reactions were prepared in FUS Assembly Buffer (5 µM GST-FUS, 1.5 mM DTT, and 0.2 U/µL RiboLock RNase Inhibitor (Thermo Scientific, EO0382)) and aggregation was initiated with TEV protease (16 µg/mL). Turbidity was measured at 395 nm in 1-minute cycles at 25°C using TECAN Spark plate reader. Parallel reactions were prepared for use in differential interference contrast (DIC) microscopy and sedimentation analysis. Once turbidity curves reached a plateau, aggregates were confirmed by DIC imaging then unlabeled RNAs were added. To account for differences in RNA length, equal mass concentration was used at 100 ng/µL, which ranged from ∼9.85 µM for the longest RNA (PSD-95 GQ2*2) to ∼14.91 µM for the shortest RNA (PSD-95) tested. Turbidity measurements were continued to monitor FUS disaggregation over time. Turbidity curves and AUC were proccessed and quantified as described above. DIC images were acquired approximately 90-100 minutes after RNA addition and sedimentation analysis was conducted 120 minutes after RNA addition. For sedimentation analysis, aggregation reactions were centrifuged at 21,000 g for 10 minutes at 4°C. Supernatant and pellet fractions were divided, then equal volumes of each fraction were analyzed by sodium dodecyl sulfate polyacrylamide gel electrophoresis (SDS-PAGE) followed by Coomassie Brilliant Blue staining. The relative abundance of FUS in each fraction was quantified using Fiji^35^ software.

## Anisotropy

MBP-FUS in TAB-K^+^ or TAB-Na^+^ was centrifuged as described above and concentration was measured with Bradford Assay. Serial dilutions of MBP-FUS were prepared, then fluorescein-tagged RNA was added to a final concentration of 8 nM. The fluorescence anisotropy of the protein-RNA mixture was measured at 25°C using TECAN Spark plate reader. At least three independent experiments were performed and measurements were plotted and fitted to the following equation:

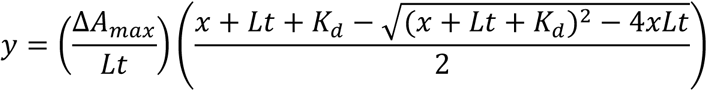

Where *ΔA_max_* is the baseline-subtracted anisotropy value, *Lt* is the total ligand concentration in nanomolar, and *K_d_* is the dissociation constant in nanomolar.

## Circular Dichroism

RNAs were diluted to 5 µM in 10 mM Tris-HCl pH 7.4 (“No Salt”) supplemented with 100 mM KCl (“K^+^”), 100 mM NaCl (“Na^+^”), or 3 mM MgCl_2_ (“Mg^2+^”). Samples were folded by heating at 90°C for 3 minutes then gradually cooled to room temperature before analysis. Folded samples were transferred to a quartz cell and spectra were analyzed at 25°C in a J-810 Spectropolarimeter (JASCO). The following parameters were used to obtain each spectra: Sensitivity (100 mdeg), Wavelength start to end (340 nm to 200 nm), Pitch (1 nm), Scanning Mode (Continuous), Scan Speed (100 nm/min), Response (0.5 s), Bandwidth (1 nm), and Accumulation (3). Spectra were smoothed using the Means-Movement method with a convolution width of 15. Respective buffer spectra were subtracted from each curve before plotting. All elipticities shown are direct readings from 5 µM RNA and not normalized by molar concentration. For melt curve analysis, sample spectra at 25°C were obtained to confirm G-Quadruplex structure. Once validated, temperature melt was conducted at 265 nm using the following parameters: Temperature start to end (15°C to 85°C), Pitch (1°C), Delay Time (2 s), Temperature Slope (3°C/min), Sensitivity (100 mdeg), Response (2 s), and Bandwidth (1 nm). Resulting spectra were smoothed using the Means-Movement method before plotting. Melt curves were fitted with the following equation:

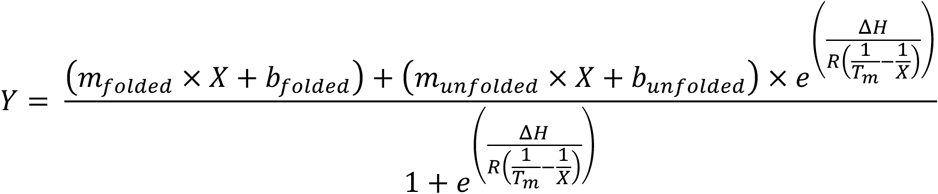

Where *m* is the slope of the baseline, *b* is the intercept of the baseline, *T_m_* is the melting temperature, *τιH* is the enthalpy change, and *R* is the ideal gas constant.

## Cell Culture

HEK 293 cells (American Type Culture Collection, “ATCC”, CRL-1573) were cultured in Dulbecco’s Modified Eagle’s Medium (DMEM; Corning, 10-013-CV) supplemented with 1% penicillin-streptomycin (Gibco, 15140-122) and 10% heat-inactivated fetal bovine serum (Cytiva, SH30396.03). Cells were plated onto poly-D-lysine-coated coverslips overnight in DMEM without antibiotic supplementation. Cells were stressed with 500 µM sodium arsenite (Alfa Aesar, 41533) for 1 hour to induce FUS phase separation and stress granule recruitment. Cells were washed with ice-cold PBS (Gibco, 10010-023) then incubated on-ice in Permeabilization Buffer (20 mM HEPES-KOH pH 7.4, 50 mM NaCl, 3 mM MgCl_2_, 300 mM Sucrose, and 0.5% Triton X-100) for 8 minutes. Once permeabilized, RNA diluted in PBS and RNasin Plus Ribonuclease Inhibitor (Promega, N2611) were added and incubated with permeabilized cells at room temperature for 30 minutes. Cells were immediately fixed using 4% paraformaldehyde (Sigma-Aldrich, 158127), then washed with PBS supplemented with 0.02% Tween 20 (BIO-RAD, 1706531) (PBS-T). Fixed cells were blocked in 3% BSA (Cell Signaling Technology, 9998S) diluted in PBS-T for 30 minutes at room temperature followed by primary staining with mouse anti-TIAR (BD Transduction Labs, 610352, 1:400) and rabbit anti-FUS (Bethyl Labs, A300-294A, 1:1000) antibodies diluted in 3% BSA overnight at 4°C. Cells were washed with PBS-T then stained with donkey anti-mouse-AF594 (Invitrogen, A-21203, 1:400) and goat anti-rabbit-AF647 (Invitrogen, A-21245, 1:400) secondary antibodies diluted in 3% BSA for 30 minutes at room temperature. Coverslips were washed with PBS-T prior to mounting to glass slides with Vectashield Antifade Mounting Medium with DAPI (Vector Laboratories Inc, H-1200). Images were acquired using 40x/1.30 oil objective on a DMi8 microscope (Leica, Germany). For each condition, 25 images were acquired in pre-defined regions across the entire coverslip, and FUS foci correlation with TIAR foci was quantified in CellProfiler^36^. Briefly, FUS and TIAR channels were enhanced for speckles (feature size = 20) and thresholds applied to select foci. DAPI channel was used to select nuclei, which were shrunk by 5 pixels to prevent under-representing peri-nuclear foci. Shrunken nuclei were inversely-masked onto FUS and TIAR channels to create a mask of cytoplasmic FUS or TIAR foci. Colocalization of the cytoplasmic FUS foci and cytoplasmic TIAR foci was measured with Pearson correlation. Imaging and quantification analyses were conducted under blinded conditions and at least three independent experiments were completed.

## Microinjection

HEK 293T cells (ATCC, CRL-3216) were cultured in DMEM (Thermo Fisher Scientific, 11995065) supplemented with 10% fetal bovine serum (Fisherbrand, FB12999102) and 1% penicillin-streptomycin (Thermo Fisher Scientific, 15140122). Approximately 1 million cells were plated into a 100-mm plastic dish, and maintained at 37°C, with 5% CO_2_ and 100% relative humidity. 48 hours before the microinjection experiments, a total of 200,000 HEK 293T cells were seeded onto a 35-mm glass bottom dish (Cellvis, D35-20-1.5-N) treated with Matrigel Matrix (Corning Life Sciences, 47743-715-EA) for 1 hour under 37 °C. 24 hours after seeding the cells, a plasmid encoding FUS-WT-eGFP (Addgene, #60362) was transfected with Lipofectamine 3000 Reagent according to manufacturer protocol (Thermo Fisher Scientific, L3000015). Immediately prior to the microinjection experiments, the cell culture medium was replaced with Extracellular Imaging Buffer (140 mM NaCl, 5 mM KCl, 10 mM HEPES, 10 mM glucose, 2 mM MgCl_2_, and 2 mM CaCl_2_, at a pH of 7.4; all chemicals were ordered from Sigma-Aldrich).

The microinjection experiments were performed on a Ti2-A inverted fluorescence microscope (Nikon, Japan) equipped with an ORCA-Fusion camera (Flash4.0 V3, Hamamatsu), a motorized stage and two motorized four-axis micromanipulators (PatchPro-5000, Scientifica). Micropipettes used in this experiment were made from borosilicate glass (World Precision Instruments) with a pipette puller (PC-10, Narishige) and filled with the RNA containing Intracellular Solution (126 mM K-gluconate, 4 mM KCl, 10 mM HEPES, 2 mM Mg-ATP, 0.3 mM Na_2_-guanosine 5′-triphosphate, 10 mM phosphocreatine, final pH 7.2, osmolarity of 270 to 290 mosmol; all chemicals were ordered form Sigma-Aldrich). The RNA species and concentration are given in the corresponding figure captions. The micropipette was connected to an Axon 700B amplifier (Molecular Devices) and to a homemade pressure recording device using an Arduino (UNO R3, ELEGOO) controlled pressure sensor (B07N8SX347, FTVOGUE, resolution of 100 Pa). Under voltage-clamp configuration, the pipette (5-10 MΩ) was inserted into a dish of HEK 293T cells expressing FUS-GFP condensates. The pipette was under a slight ejection pressure (amplitude ∼ 1 kPa) to prevent clogging. Then a negative suction pressure was applied to form a giga-seal near the target condensate. Switching to current-clamp mode (with I = 0), the voltage signal was recorded, and pulses of suction pressure (amplitude > 10 kPa) were applied to rupture the membrane to achieve a whole-cell patch-clamp configuration. After which an ejection pressure (∼ 10 kPa) was applied through the patch pipette. The membrane potential recordings were processed through a 2-kHz filter, digitized at a rate of 10 kHz, and acquired using Clampex 10.2 software (Molecular Devices). The condensate-containing cells were imaged in GFP channel and Cy5 channel to evaluate the effect of RNA injection.

## Single-Molecule FRET

For smFRET measurements, the details of instrumentation and PEGylated slide preparation were as described^37,38^. Briefly, the microfluidic sample chamber was created between the plasma cleaned slide and the coverslip coated with polyethylene glycol (PEG) and biotin-PEG. Annealed RNA molecules were immobilized on the PEG-passivated surface via biotin-neutravidin interaction. All smFRET measurements were carried out in imaging buffer containing an oxygen scavenger system to stabilize fluorophores (10 mM Tris-HCl, pH 7.5, 100 mM KCl, 10 mM trolox, 0.5% (w/v) glucose, 1 mg/mL glucose oxidase and 4 g/ml catalase)^37^. All smFRET assays were performed at room temperature (∼23°C±2°C). Wide-field prism-type total internal reflection fluorescence (TIRF) microscopy was used with a solid-state 532nm diode laser to generate an evanescent field of illumination to excite the fluorophores (Cy3 or Cy5) at the sample chamber. Fluorescence signals from Cy3 (donor) and Cy5 (acceptor) were simultaneously collected using a water immersion objective and sent to a charge-coupled device (CCD) camera after passing through the dichroic mirror (cut off = 630 nm). Movies were recorded over different regions of the imaging surface with a time resolution of 100ms as a stream of imaging frames. FRET histograms were built by collecting FRET values from over 5000 molecules in 20 different fields of view (21 frames of 20 short movies). Long movies (1200 frames, i.e., 120 s) were recorded to look through the molecular behavior using MATLAB script.

## Bioinformatic Analysis

The impact of liquid-liquid phase separation (LLPS) on the FUS interactome was analyzed using publicly available RNA-seq data from EMBL-EBI (Accession E-MTAB-8456)^39^. QC, alignment, and conversion of raw data to featurecounts were done using TrimGalore (0.6.10) (https://zenodo.org/records/7598955), STAR (2.7.10b)^40^, and Rsubread (2.8.2)^41^ respectively. All statistical analyses were performed using R Statistical Software (v4.1.3) (https://www.R-project.org/). Aligned reads were then compiled into a single bedfile for each experimental group using BEDOPS toolkit (2.4.41)^42^.

G-quadruplex (G4) structures were analyzed using data from rG4-Seq experiments downloaded from NCBI GEO (Accession GSE77282)^43^. The liftover of G4 genomic coordinates (rG4-Seq) from hg19 to hg38 genome assembly was performed using UCSC LiftOver (377)^44^. Overlaps between G4 and previously mentioned FUS interactome datasets were found using the intersect function of bedtools (v.2.31.1)^45^. Following the methods of a previously published G4-analysis, the G4 structures were stratified into three equally sized groups according to their stability score, defined as a fraction of the reads with reverse transcription stops, and here named “G4 Probability”^46^. The three groups are labeled “low”, “medium”, and “high”. To quantify enrichment between soluble FUS interactome and droplet FUS interactome, we first selected sequences classified as “high G4 probability” shared between the two groups. We then calculated an enrichment score for each shared sequence as the ratio of its occurrence in the soluble FUS interactome to its occurrence in the droplet FUS interactome.

Publicly available datasets for CLIP-seq of TAF15^47^ and FUS^48^ were obtained via NCBI GEO (GSE43294 and GSE40651 respectively). Data were converted from hg18 to hg38 using UCSC LiftOver. Regions were then overlapped with rG4seq using the intersect function of bedtools.

To evaluate the specificity for overlaps of G4 regions and soluble phase or droplet phase, permutation testing was performed in both directions with bedtools shuffle for 100 randomized versions of either G4 regions or non-G4 regions within hg38. The results were then analyzed with python by comparing the counts of observed overlaps and randomized overlaps. An empirical p-value was computed and included in a histogram of the permutation results.

## Statistical Analysis

Unless otherwise stated, all statistical analyses were performed in GraphPad Prism 10 and the method used is indicated in the figure legends.

### RESULTS

## Strong RNA inhibitors prevent and reverse FUS phase separation in a concentration-dependent manner

We aimed to achieve a mechanistic understanding of the different activities of the RNA inhibitors we have identified^30^, with a particular interest in understanding the superior activity of strong RNA inhibitors, such as S1. A distinct feature we observed for strong RNA inhibitors is their ability to inhibit FUS LLPS, while weak inhibitors do not. Thus, we focused on this feature and quantitatively assessed the effects of specific RNA inhibitors on FUS LLPS in the absence of aggregation (**Fig. 1A**). We used a FUS LLPS assay in which TEV protease cleaves the MBP tag from MBP-TEV-FUS, triggering formation of FUS droplets within 60 minutes that subsequently coalesce into larger, liquid-like structures (**Fig. 1B**). Imaging analysis pipelines were developed to generate phase diagrams that quantitatively map FUS phase behavior by showing droplet number and FUS enrichment over time (**Fig. 1C**). Without RNA, FUS enriches in droplets as their number decreases and size increases over time (**Figs. 1B-C, S1A**). Excess RNA S1 can completely inhibit FUS LLPS, as previously reported^30^ (**Fig. 1C-D**). Interestingly, low concentration of RNA S1 leads to smaller droplets (**Figs. 1D and S1A**; 1 µM) with less FUS enriched, but the number of droplets increases compared to control (**Fig. 1C-D**; 1 µM), suggesting RNA S1 inhibits FUS LLPS by both inhibiting its recruitment into droplets and droplet coalescence. Further increasing the concentration of RNA S1 to 32 µM leads to decreases in FUS droplet size (**Fig. S1A**), number, and enrichment score across each time point, thus completely preventing its phase separation (**Fig. 1C-D**). These concentration-dependent effects of RNA S1 on FUS LLPS are further corroborated by turbidity assay, a higher-throughput alternative to imaging analysis. At low RNA concentrations (1, 2, or 4 µM), the turbidity (**Fig. 1E)** and the corresponding area under the curve (AUC) (**Fig. 1F**) are increased relative to control, reflecting the increased droplet number (**Fig. 1D**) that diffracted more light. Higher RNA S1 concentrations reduce turbidity, reflecting fewer and smaller droplets. When RNA S1 concentration is increased to 16 and 32 µM, FUS LLPS is fully prevented (**Fig. 1E-F**). Another strong RNA inhibitor, RNA S2^30^, exhibits similar concentration-dependent inhibition of FUS LLPS, as observed in droplet images (**Fig. S1B**) and turbidity changes (**Fig. S1C-D**). In contrast, RNA W1, a representative weak inhibitor previously shown to reduce turbidity caused by FUS aggregation by 50%^30^, has minimal activity in preventing FUS phase separation (**Figs. 1G-H, S1E**). In turbidity assays, RNA W1 slightly increased turbidity at all tested concentrations; however, this effect was not concentration-dependent (**Fig. 1I-J**). Thus, strong RNA inhibitors suppress FUS LLPS in a concentration-dependent manner, whereas weak inhibitors do not.

An intriguing feature of strong RNA inhibitors was their ability to reverse FUS aggregation^30^, so we tested whether they also reverse FUS phase separation. Adding low-concentration RNA S1 to pre-formed droplets reduces the droplet size and FUS enrichment while increasing droplet number (**Figs. 1K-L, S1F**; 1 µM S1), suggesting RNA S1 may be breaking apart pre-formed droplets as an initial step of reversing, which was directly observed for larger droplets over time (**Fig. S1G**). Higher RNA S1 levels further decreased all measured parameters, and at 32 µM, led to droplet dissolution (**Figs. 1K-L, S1F**). Turbidity experiments report a similar dose-dependent reduction in FUS turbidity by RNA S1 (**Fig. 1M-N**). Similarly, RNA S2 drastically reverses FUS phase separation in a dose-dependent manner in both imaging and turbidity assay (**Fig. S1H-J**). In contrast, RNA W1 shows only mild reversal of FUS LLPS, with increased droplet number, reduced size, and FUS enrichment (**Figs. 1O-P, S1K**). Unlike S1 or S2, W1 shows no concentration-dependent effect and fails to fully reverse phase separation. Turbidity assays confirm this, showing an “enhanced” turbidity curve, reflecting increased droplet number without further changes at higher W1 concentrations (**Fig. 1Q-R**).

Encouraged by the strong activity of RNA S1, we asked whether it could also reverse FUS phase separation in a cellular environment. To evaluate this effect in real-time, we utilized microinjection^49–51^ to directly introduce RNA S1 into HEK 293T cells overexpressing FUS-eGFP, which is prone to forming condensates even without additional stressors^16^. We first confirmed that RNA S1 localizes to pre-formed FUS-eGFP condensates by injecting a low concentration (0.5 µM) of Cy5-S1 into the cells, which rapidly diffused from the injection site at the top of the cell to cytoplasmic FUS-eGFP condensates C1 and C2 (**Fig. S1L**, marked by yellow and white arrows). We then tested the disassembly activity of RNA S1 at a higher concentration (100 µM) (**Fig. 1S**). During pre-injection, the liquid-like property of the condensate (C2, red arrows) was confirmed by shape change caused by a brief negative pressure that sucked the condensate into the pipette (**Fig. 1S**; panels 1-3). Upon injecting RNA S1 (**Fig. 1S**; panels 4-8), condensates C2 and C3, which were nearest to the injection site, were dissolved within 1 minute. Thus, strong RNA inhibitors, such as RNA S1, can effectively reverse FUS LLPS in cells.

## rG4 is the specific structural determinant for RNA S1 activity in mitigating FUS LLPS

Given the therapeutic potential of strong RNA inhibitors and the fact that RNAs S1 and W1 are similar in length (**Table S1**) yet exhibit distinct capacities to modulate FUS LLPS, we aimed to uncover structural determinants of strong RNA inhibitors. Our previous work indicated that the compact and rigid structure of strong RNA inhibitors hinders the multivalent binding of FUS, thus preventing its aggregation^30^. Conversely, more extended and flexible RNAs, such as the weak inhibitor U50, facilitate multivalent interactions, leading to multimer complex formation and LLPS^30^. Thus, we hypothesized that promoting the formation of compact and rigid structures in W1 could enhance its ability to mitigate FUS LLPS. Mg^2+^ is effective at inducing RNA folding and stabilizing RNA tertiary structures^52,53^. Indeed, using single-molecule Förster resonance energy transfer (smFRET) (**Fig. S2A**), we confirmed the structural compaction and stabilization of RNA W1 in the presence of Mg^2+^ (**Fig. S2B**), evidenced by higher FRET efficiency and narrower distribution. However, this structural change did not improve W1’s activity in reversing FUS LLPS (**Fig. S2C-D**). More surprisingly, the activity of strong RNA inhibitors S1 and S2 is sensitive to the presence of Mg^2+^. smFRET measurements show that Mg^2+^ also induces a more compact and rigid structure in RNA S1 (**Fig. S2E**). However, RNA S1 activity in reversing FUS LLPS is completely lost in Mg^2+^ (**Fig. S2F-G**; “+Mg^2+^”), despite maintaining recruitment to FUS droplets (**Fig. S2H**). Similar sensitivity to Mg^2+^ is also observed for RNA S2 (**Fig. S2I-K**). These results suggest that a general compact structure does not define strong RNA activity and that Mg^2+^ may alter an important structure that is critical for RNA S1 and S2’s activity.

In order to identify the structural determinant of strong RNA inhibitors, we employed circular dichroism (CD). RNA S1 exhibits a main positive peak at ∼265 nm, along with a negative peak at ∼240 nm, indicative of parallel RNA G-quadruplex (rG4) structure^54–56^, which is enhanced in the presence of K^+^ compared to Na^+^ (**Fig. 2A**) and consistent with QGRS Mapper^31^ prediction (**Fig. 2B**, **Table S1**). Moreover, thermal stability analysis confirmed the rG4 structure in S1 is more stable in K^+^ (*T_m_* = 50.64°C) than in Na^+^ (*T_m_* = 36.98°C) (**Fig. 2C**). The rG4 structure is disrupted in the presence of Mg^2+^, as indicated by a lower 265 nm peak (**Fig. 2A**; purple trace). In contrast, both CD and QGRS Mapper indicate that RNA W1 does not form rG4 (**Fig. S2L, Table S1**).

**Figure 2.**
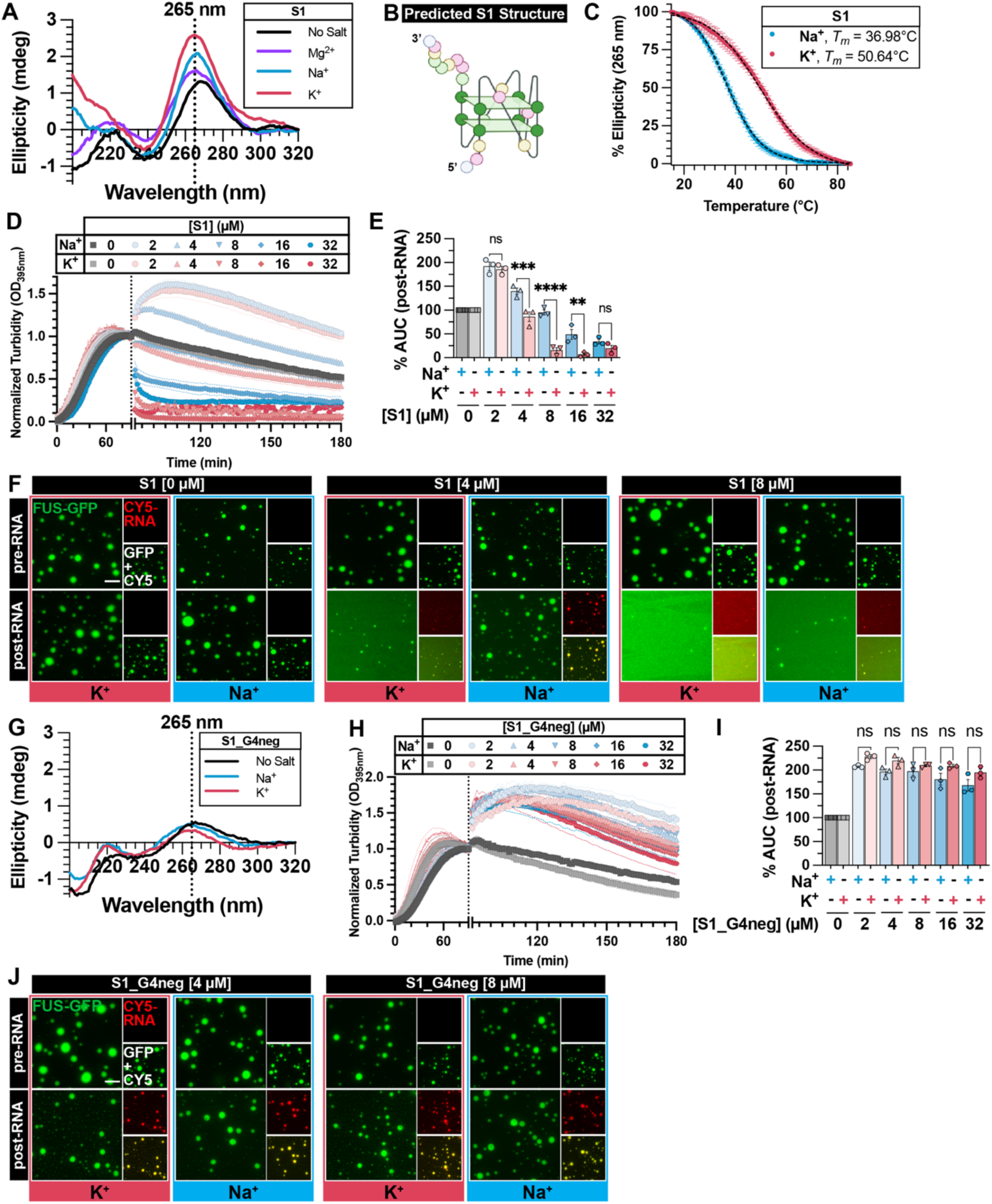
rG4 is the specific structural determinant for RNA S1 activity in mitigating FUS LLPS. (**A, G**) Representative circular dichroism (CD) spectra of RNA S1 (**A**) or S1_G4neg (**G**) collected at 25°C in indicated buffer condition. (**B**) Schematic representation of predicted rG4 formed by RNA S1. RNA structures were created with Biorender.com. (**C**) CD thermal melt curves of RNA S1 in Na^+^ (blue) or K^+^ (red). CD signals were collected at 265 nm from 15°C to 85°C. Data plotted represent mean±SEM (n=3). Black dashed lines represent the fitted curve used to obtain *T_m_*. (**D, H**) Turbidity measurement of FUS LLPS reversal with RNA S1 (**D**) or S1_G4neg (**H**) in TAB-Na^+^ (blue traces) or TAB-K^+^ (red traces). LLPS of FUS (2 µM) was initiated, and monitored by turbidity at 395 nm. After 90 min, data collection was paused to add RNA. Increasing concentration of RNA is indicated by a more saturated color. Data represent mean±SEM (n=3). The 0 µM RNA control in TAB-K^+^ plotted in Fig. 2D and Fig. S2F are the same because these experiments were conducted simultaneously. (**E, I**) AUC was calculated from turbidity curves in Fig. 2D (**E**) or Fig. 2H (**I**) after RNA was added. Data represent mean±SEM (n=3). Ordinary one-way ANOVA with Šidák’s multiple comparisons test was used to compare the means between TAB-Na^+^ and TAB-K^+^ at each RNA concentration (ns P>0.05, **P≤0.005, ***P≤0.0005, ****P<0.0001). (**F, J**) Representative images of FUS droplets before (top row) and after (bottom row) the addition of RNA S1 (**F**), or S1_G4neg (**J**). LLPS of FUS (1.8 µM FUS supplemented with 0.2 µM FUS-GFP) was monitored by droplet imaging before and 30 minutes after the addition of RNA (4 or 8 µM unlabeled RNA supplemented with 75 nM Cy5-RNA). Scale bar is 5 µm.

To determine whether rG4 is the specific structural determinant of RNA S1’s activity in mitigating FUS LLPS, turbidity experiments were repeated in the presence of rG4-stabilizing ion K^+^ and compared to results obtained in the presence of Na^+^, where rG4 is less stable (**Fig. 2D-E**). In the following experiments, RNAs are melted and refolded in the respective buffers to ensure proper structure. While RNA S1 can reverse FUS phase separation in a concentration-dependent manner in both buffer conditions, its activity is significantly stronger in K^+^ (**Fig. 2D-F**). The observed difference in activity is not a result of different binding affinity in K^+^ or Na^+^, as S1 has similar binding affinity for FUS (**Fig. S2M**) and localizes to pre-formed FUS droplets (**Fig. 2F**) in both buffer systems.

To further confirm the contribution of rG4 structure to RNA S1’s activity, we scrambled the sequence of RNA S1 to prevent rG4 structure (S1_G4neg), which was confirmed by CD (**Fig. 2G**). As a result, S1_G4neg completely loses its ability to reverse FUS phase separation in both K^+^ and Na^+^ conditions (**Fig. 2H-J**), and behaves similarly to weak inhibitor W1, which is both salt- and concentration-insensitive (**Fig. S2N-O**). Despite a disrupted structure, S1_G4neg maintains its localization to FUS droplets (**Fig. 2J**) and strong binding affinity to FUS (**Fig. S2P**) in both buffers. Collectively, these results establish rG4 as the principal structural element governing RNA S1’s ability to reverse FUS phase separation.

## rG4, not hairpin structure, also determines RNA S2 activity in mitigating FUS LLPS

To determine whether the contribution of rG4 in reversing FUS phase separation is unique to RNA S1, and whether other compact structures are capable of mitigating FUS LLPS, we examined RNA S2. RNA S2 is predicted to be able to adopt both an rG4 and a stem-loop structure (**Fig. S3A, Table S1**), both of which are inherently compact, which would explain the little change observed in smFRET with/without Mg^2+^ (**Fig. S2I)** and allows us to dissect the contributions of these distinct compact structural elements to its activity. CD spectra confirmed RNA S2 forms rG4 in K^+^, which is less stable in Na^+^ (**Fig. S3B**), correlating with stronger reversal activity against FUS LLPS in K^+^ compared to Na^+^ (**Fig. S3C-E**) without altering localization to FUS droplets or binding affinity (**Fig. S3E-F**). To further delineate rG4 vs. stem-loop contributions, we designed RNA S2_G4neg, a minimally scrambled mutant that disrupts rG4 but retains the stem-loop (**Fig. S3G, Table S1**). CD confirms that RNA S2_G4neg does not adopt rG4 structure in K^+^ or Na^+^ (**Fig. S3H**). Consistently, RNA S2_G4neg failed to reverse FUS phase separation, and similar to other weak inhibitors, does not show enhanced activity in K^+^ compared to Na^+^ (**Fig. S3I-K**). Despite lost activities, RNA S2_G4neg localizes to FUS droplets and maintains a strong affinity for FUS, like RNA S2 (**Fig. S3K-L**). These results establish rG4 as a key structural element, that is distinct from other compact structures, in driving RNA’s ability to reverse FUS LLPS.

## FUS-binding rG4 is a defining criterion for identifying strong RNA inhibitors of FUS LLPS and aggregation

Our results suggest that rG4 structure is a common feature amongst strong RNA inhibitors and could potentially be used to identify RNA inhibitors of FUS LLPS. Thus, we utilized QGRS-mapper^31^ to predict the G4-forming propensity of other RNA inhibitors of FUS aggregation identified in our previous study^30^ (**Table S2**). This includes 8 strong inhibitors (S1-S8), 9 weak inhibitors (W1-W9), and 6 RNAs with no activity (N1-N6). Notably, over 60% of the strong inhibitors were predicted to contain rG4 elements, while none of the weak inhibitors or no activity RNAs were (**Table S2**). This suggests that rG4 may be a general property of the strong RNA inhibitors capable of mitigating FUS LLPS. Thus, we hypothesized that other rG4-forming RNAs that interact with FUS would exhibit strong RNA activity. We began by assessing TERRA, which is an archetypal rG4 that binds FUS with nanomolar affinity^57^. CD spectra show that, unlike other rG4 tested thus far, TERRA folded in K^+^ or Na^+^ displays a similarly strong rG4 signature (**Fig. S4A**), indicating equally-stabilized rG4 structures in both buffers. Accordingly, TERRA can strongly reverse FUS LLPS in both K^+^ and Na^+^ buffers (**Fig. 3A-C**).

**Figure 3.**
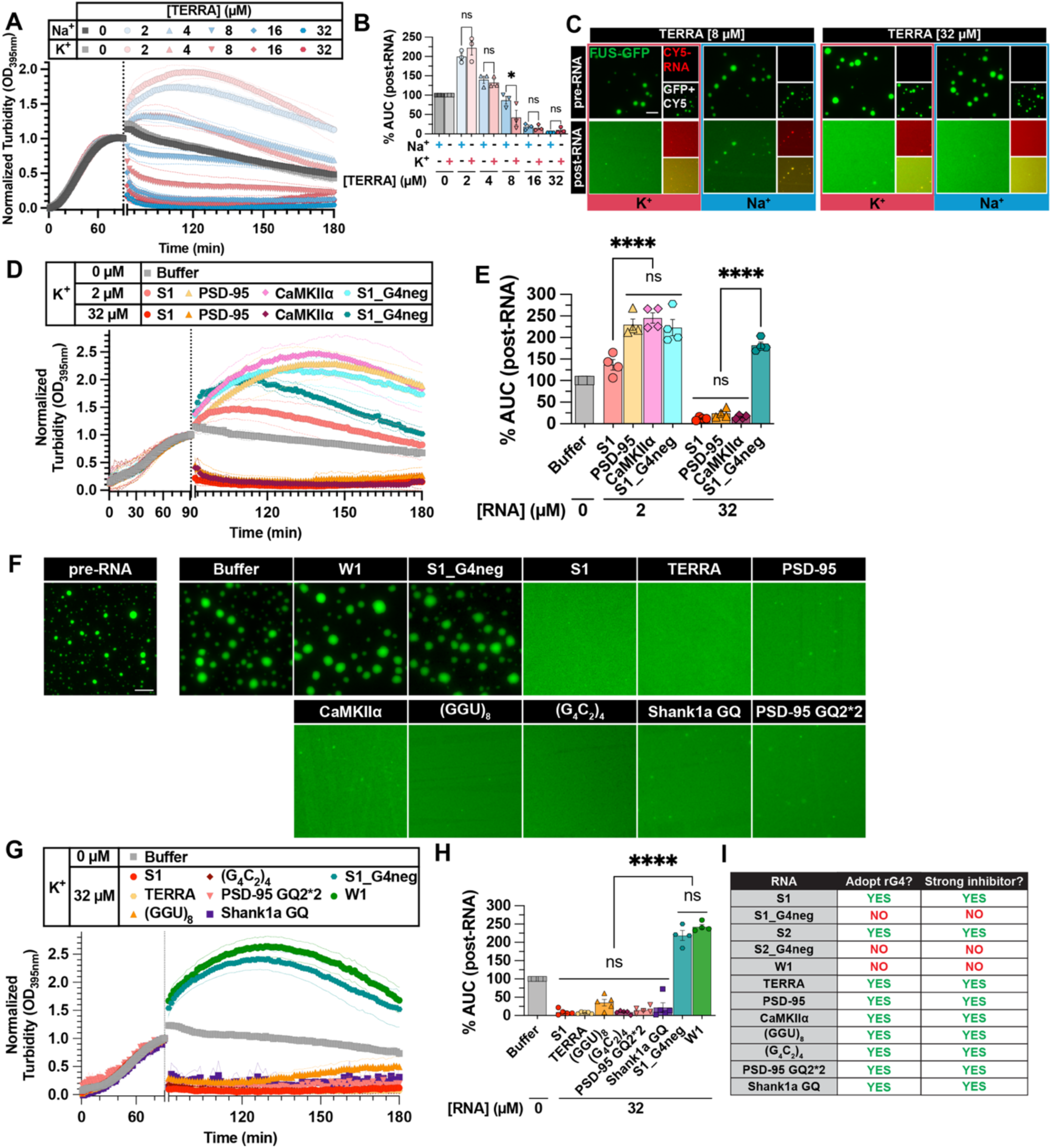
FUS-binding rG4 is a defining criterion for identifying strong RNA inhibitors of FUS LLPS and aggregation. **(A)** Turbidity measurement of FUS LLPS reversal with TERRA in TAB-Na^+^ (blue traces) or TAB-K^+^ (red traces). LLPS of FUS (2 µM) was initiated, and monitored by turbidity at 395 nm. After 90 min, data collection was paused to add RNA. Increasing concentration of RNA is indicated by a more saturated color. Data represent mean±SEM (n=3). **(B)** AUC was calculated from turbidity curves in Fig. 3A after RNA was added. Data represent mean±SEM (n=3). Ordinary one-way ANOVA with Šidák’s multiple comparisons test was used to compare the means between TAB-Na^+^ and TAB-K^+^ at each RNA concentration (ns P>0.05, *P≤0.05). **(C)** Representative images of FUS droplets before and after the addition of TERRA. LLPS of FUS (1.8 µM FUS supplemented with 0.2 µM FUS-GFP) was monitored by droplet imaging before and 30 minutes after addition of TERRA (8 or 32 µM unlabeled RNA supplemented with 75 nM Cy5-RNA). Scale bar is 5 µm. **(D)** Turbidity measurement of FUS LLPS reversal with indicated RNA in TAB-K^+^. LLPS of FUS (2 µM) was initiated, and monitored by turbidity at 395 nm. After 90 min, data collection was paused to add RNA (2 µM or 32 µM). Data represent mean±SEM (n=4). **(E)** AUC was calculated from turbidity curves in Fig. 3D after RNA was added. Data represent mean±SEM (n=4). Ordinary one-way ANOVA with Šidák’s multiple comparisons test was used to compare the means between RNAs at each concentration (ns P>0.05, ****P<0.0001). **(F)** Representative images of FUS droplets before (“pre-RNA”) and after the addition of indicated RNAs. LLPS of FUS (1.8 µM FUS supplemented with 0.2 µM FUS-GFP) was monitored by droplet imaging before and 60 minutes after the addition of 32 µM unlabeled RNA. Scale bar is 5 µm. **(G)** Turbidity measurement of FUS LLPS reversal with indicated RNA in TAB-K^+^. LLPS of FUS (2 µM) was initiated, and monitored by turbidity at 395 nm. After 90 min, data collection was paused to add RNA (32 µM). Data represent mean±SEM (n=4-5). **(H)** AUC was calculated from turbidity curves in Fig. 3G after RNA was added. Data represent mean±SEM (n=4-5). Ordinary one-way ANOVA with Tukey’s multiple comparisons test was used to compare the means between RNAs at each concentration (ns P>0.05, ****P<0.0001). **(I)** Summary table of RNAs tested, indicating whether parallel rG4 was observed by CD (“Adopt rG4”) and whether the RNA displayed characteristics of strong RNA inhibitors by turbidity measurements (“Strong inhibitor”).

To further support these results, we then tested more FUS-binding rG4. Recently, Ishiguro et al. reported two rG4-forming sequences from the 3’-UTRs of mRNAs encoding PSD-95 and CaMKIIα that bind FUS^58,59^. Although enhanced FUS LLPS in the presence of these RNAs was reported^59^, we noted that the concentrations tested (0.5 µM - 8 µM) were in a range where we also observed enhanced FUS turbidity with strong inhibitors RNAs S1 and S2 (**Figs. 2D-E, S3C-D**). Therefore, we tested whether these RNAs may be able to reverse FUS LLPS at higher concentrations in rG4-stabilizing buffer conditions. CD confirmed PSD-95 and CaMKIIα adopt rG4 in K^+^ buffer (**Fig. S4B-C**). Turbidity experiments show that, at low concentrations (2 µM), PSD-95 and CaMKIIα cause enhanced FUS turbidity (**Fig. 3D-E**), consistent with the previous report^59^. However, similar to RNA S1, both PSD-95 and CaMKIIα showed a concentration-dependent activity and completely reversed FUS LLPS at high concentration (32 µM), while S1_G4neg cannot (**Fig. 3D-F**).

To test this idea further, we examined another panel of rG4 comprised of known FUS-binding motifs or known to interact with FUS (**Table S1**). This rG4 panel includes (GGU)_8_, which is composed of a putative FUS-binding sequence^60–62^, the C9orf72 hexanucleotide repeat sequence (G_4_C_2_)_463,64_, and short sequences from the 3’-UTRs of mRNAs encoding Shank1 and PSD-95 (i.e., PSD-95 GQ2)^65^ (**Fig. S4D-G**). We chose to double the sequence of PSD-95 GQ2^65^ (PSD-95 GQ2*2) so that its length matches the other RNAs studied. Remarkably, all rG4 RNAs tested can fully reverse FUS LLPS (**Fig. 3F-H**).

In addition to reversing FUS LLPS, our original strong RNAs (S1 and S2) were capable of reversing FUS aggregation^30^, an activity that is critical for therapeutic application. Given their activity against FUS LLPS, we assessed whether this new panel of rG4 RNA inhibitors was also capable of mitigating FUS aggregation. Remarkably, using assay conditions that favor the formation of FUS aggregates^30^, all rG4-forming RNAs tested showed similarly strong activity in reversing FUS aggregation, whereas non-rG4-forming RNAs (S1_G4neg and W1) could not (**Fig. S4H-L**). In summary, FUS-binding rG4 provides a robust criterion for identifying short RNA oligonucleotides with strong activity to reverse FUS LLPS and aggregation (**Fig. 3I**).

## Increasing rG4 repeat numbers in RNA S1 switches it from inhibiting to nucleating FUS assembly

Our results show that FUS-binding rG4 can mitigate FUS aggregation and have the potential to be developed as therapeutics for FUS-ALS/FTD^30^; thus, we sought to further enhance the activity of RNA inhibitors by enhancing their G4-forming capability. Using RNA S1 as a template, we first tested whether increasing the repeat number of the rG4 element can enhance its activity. We began with an oligonucleotide containing two tandem repeats of S1 sequence (S1*2) and confirmed using CD that S1*2 adopts rG4 structure in K^+^ and Na^+^ (**Fig. 4A**). Similar to S1, RNA S1*2 is able to reverse FUS phase separation in a concentration-dependent manner, and it is stronger in K^+^ than in Na^+^ at lower concentrations (**Fig. 4B-C**). However, despite having an increased propensity to form rG4 in both K^+^ and Na^+^ than RNA S1 (**Fig. 4D-E**), when directly compared using the same mass concentration, RNA S1*2 does not exhibit enhanced activity compared to RNA S1 in either K^+^ or Na^+^ (**Fig. 4F-I**), suggesting that increasing the length of the rG4 element does not enhance RNA activity against FUS LLPS.

**Figure 4.**
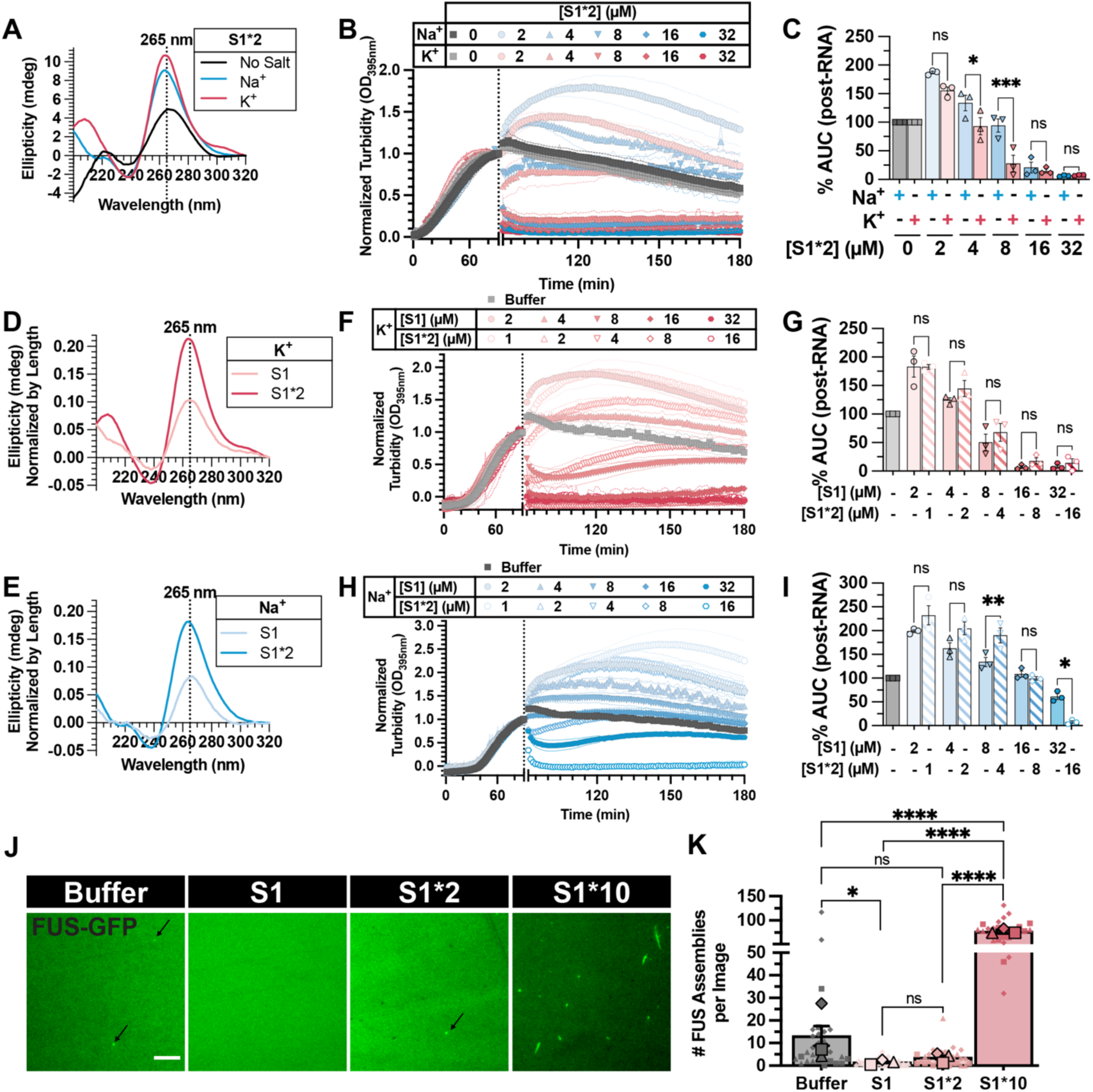
Increasing rG4 repeat numbers in RNA S1 switches it from inhibiting to nucleating FUS assembly. **(A)** Representative CD spectra of RNA S1*2 collected at 25°C in indicated buffer. **(B)** Turbidity measurement of FUS LLPS reversal with S1*2 in TAB-Na^+^ (blue traces) or TAB-K^+^ (red traces). LLPS of FUS (2 µM) was initiated, and monitored by turbidity at 395 nm. After 90 min, data collection was paused to add RNA. Increasing concentration of RNA is indicated by a more saturated color. Data represent mean±SEM (n=3). **(C)** AUC was calculated from turbidity curves in Fig. 4B after RNA was added. Data represent mean±SEM (n=3). Ordinary one-way ANOVA with Šidák’s multiple comparisons test was used to compare the means between TAB-Na^+^ and TAB-K^+^ at each RNA concentration (ns P>0.05, *P≤0.05, ***P≤0.0005). (**D, E**) CD spectra of RNA S1 (Fig. 2A) and S1*2 (Fig. 4A) were normalized by RNA length and re-plotted to compare spectra in K^+^ (**D**) and Na^+^ (**E**). (**F, H**) Turbidity measurement of FUS LLPS reversal with RNA S1 or S1*2 in TAB-K^+^ (**F**) or TAB-Na^+^ (**H**). LLPS of FUS (2 µM) was initiated, and monitored by turbidity at 395 nm. After 90 min, data collection was paused to add RNA. Increasing concentration of RNA is indicated by a more saturated color. S1 data are represented by filled symbols and S1*2 data are represented by open symbols. Data represent mean±SEM (n=3). (**G, I**) AUC was calculated from turbidity curves in Fig. 4F (**G**) and Fig. 4H (**I**) after RNA was added. Data represent mean±SEM (n=3). Ordinary one-way ANOVA with Šidák’s multiple comparisons test was used to compare the means between RNAs at comparable concentrations, accounting for the difference in length between S1 and S1*2 (ns P>0.05, *P≤0.05, **P≤0.005). **(J)** Representative images of FUS assemblies formed in the presence of RNA S1 (2 µM), S1*2 (1 µM), or S1*10 (200 nM). LLPS of FUS-GFP (500 nM) in TAB-K^+^ was initiated by adding TEV protease to cleave the MBP tag of MBP-TEV-FUS-GFP in the presence of indicated RNAs. For each trial, images were acquired 120 minutes after TEV addition (n=3). Scale bar is 5 µm and arrows point to FUS assemblies. **(K)** The average number of FUS assemblies per image was quantified from images collected in Fig. 4J. Bars represent mean±SEM of three independent experiments, large symbols represent trial means (n=3) and small datapoints represent individual image values (n=30). Ordinary one-way ANOVA with Tukey’s multiple comparisons test was used to compare the means between RNAs. (ns P>0.05, *P≤0.05, ****P<0.0001).

We reasoned that multiple FUS binding motifs on a single RNA might serve as a template for FUS oligomerization^27^, thus reducing the RNA’s activity in mitigating FUS LLPS. Therefore, we hypothesized that further increasing rG4 repeats could nucleate FUS phase separation. To test this hypothesis, we generated a ten-repeat S1 sequence (S1*10). Since the yield of *in vitro* transcription for longer RNA is not high enough for turbidity assays, a modified droplet imaging assay was developed (**Fig. S5A**) to examine the effect of S1*10 on FUS phase separation. When comparing S1 or S1*2 against S1*10 at the same mass concentration (2 µM, 1 µM, and 200 nM, respectively) (**Fig. 4J-K**) or molar concentration (200 nM) (**Fig. S5B-C**), a significant increase in the number of FUS assemblies was observed only when S1*10 is present. Surprisingly, in contrast to the droplet-like assemblies formed without RNA, when S1*10 is added, FUS forms small, fibril-like structures (**Figs. 4J, S5D**), similar to a previous report suggesting RNA acts as the template to seed FUS fibril formation^29^. These results highlight a switch-like regulatory mechanism where the repeat number of the rG4 element dictates RNA regulation of FUS phase separation: longer RNAs with multiple G4 elements promote FUS assembly, whereas shorter G4-containing RNAs inhibit it.

## Chemical modifications enhancing rG4 stability strengthen RNA activity against FUS phase separation

Since increasing RNA length reduced RNA S1’s inhibitory effect on FUS phase separation despite higher rG4 content, we explored stabilizing the rG4 without lengthening the RNA. We introduced chemical modifications, including phosphorothioate backbone and 2’-O-methylation (2’-OMe) modifications to RNA S1, generating S1^P+M^. Circular dichroism measurement showed that S1^P+M^’s rG4 was similarly stable in K⁺ and Na⁺ (**Fig. S6A**), and confirmed the modifications imparted greater thermal stability than S1 (**Fig. 5A-B**). Functionally, although S1^P+M^ did not outperform S1 in K⁺ (**Fig. 5C-D**), it exhibited significantly enhanced activity in Na⁺ (**Fig. 5E-F**). Remarkably, S1^P+M^ retained activity even in 3 mM Mg²⁺, unlike the Mg²⁺-sensitive S1 (**Figs. 5G-I, S2F-G**). These data indicate that chemical modifications stabilizing the rG4 structure can mitigate RNA inhibitor sensitivity to ionic conditions.

**Figure 5.**
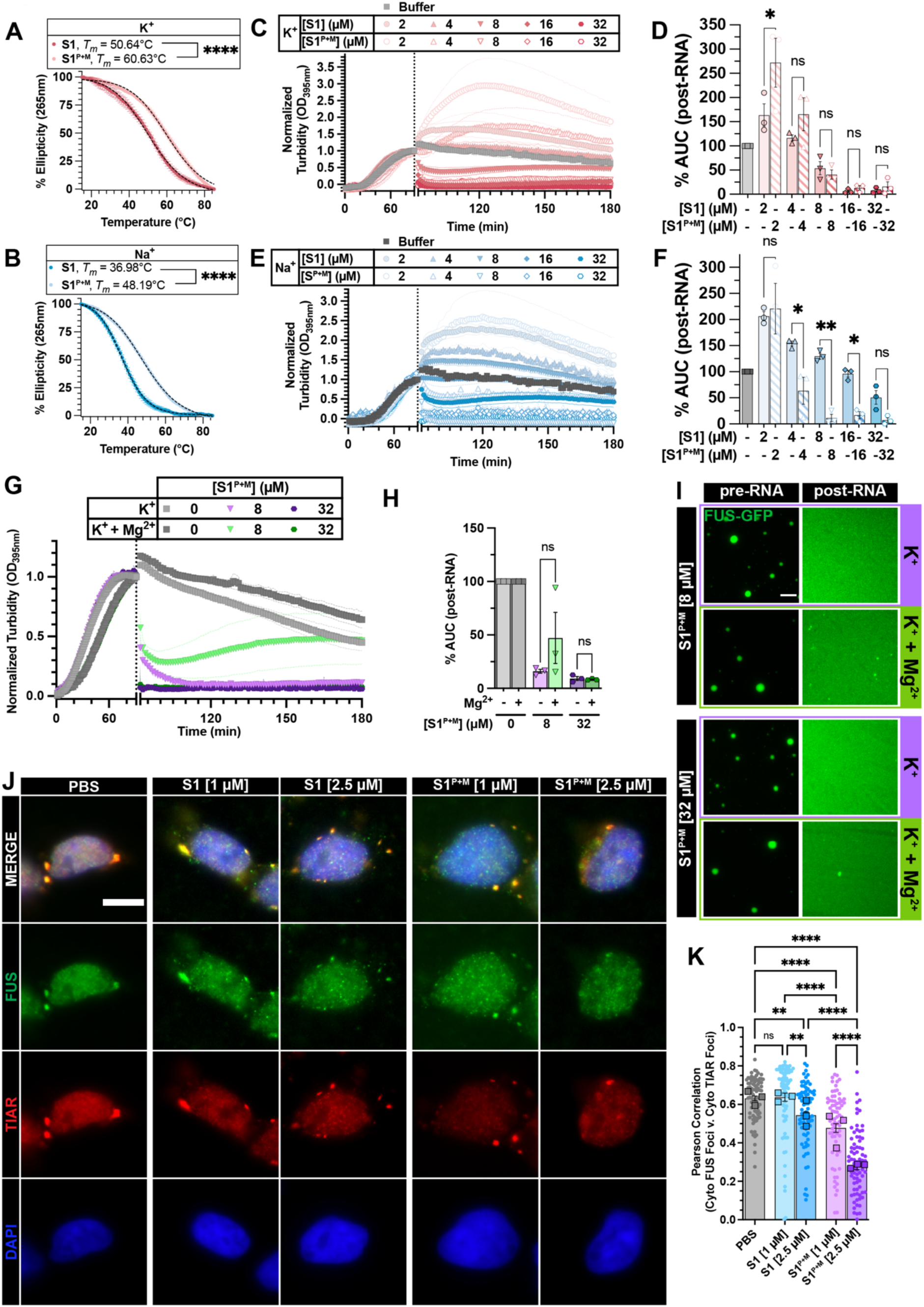
Chemical modifications enhancing rG4 stability strengthen RNA activity against FUS phase separation. (**A, B**) CD thermal melt curves of RNA S1 and S1^P+M^ in K^+^ (**A**) or Na^+^ (**B**). Data plotted represent mean±SEM (n=3). Black dashed lines represent the fitted curve used to obtain *T_m_*. Thermal melt curves of RNA S1 are replotted from Fig. 2C in order to compare with RNA S1^P+M^. Extra sum-of-squares F Test was used to compare *T_m_* of S1 and S1^P+M^ in each buffer condition. (****P<0.0001). (**C, E**) Turbidity measurement of FUS LLPS reversal with RNA S1 or S1^P+M^ in TAB-K^+^ (**C**) or TAB-Na^+^ (**E**). LLPS of FUS (2 µM) was initiated, and monitored by turbidity at 395 nm. After 90 min, data collection was paused to add RNA. Increasing concentration of RNA is indicated by a more saturated color. S1 data are represented by filled symbols, and S1^P+M^ data are represented by open symbols. Data represent mean±SEM (n=3). (**D, F**) AUC was calculated from turbidity curves in Fig. 5C (**D**) and Fig. 5E (**F**) after RNA was added. Data represent mean±SEM (n=3). Ordinary one-way ANOVA with Šidák’s multiple comparisons test was used to compare the means between S1 and S1^P+M^ at each concentration (ns P>0.05, *P≤0.05, **P≤0.005). **(G)** Turbidity measurement of FUS LLPS reversal with S1^P+M^ in TAB-K^+^ ± Mg^2+^. FUS (2 µM) LLPS was initiated, and monitored by turbidity at 395 nm. After 90 min, data collection was paused to add S1^P+M^ diluted in 10 mM Tris-HCl pH 7.4 supplemented with 50 mM KCl (purple symbols) or supplemented with 50 mM KCl + 3 mM MgCl_2_ (green symbols). Data represent mean±SEM (n=3). **(H)** AUC was calculated from turbidity curves in Fig. 5G after RNA was added. Data represent mean±SEM (n=3). Ordinary one-way ANOVA with Šidák’s multiple comparisons test was used to compare the means (ns P>0.05). **(I)** Representative images of FUS droplets in TAB-K^+^ ± 3 mM MgCl_2_ before and after the addition of RNA S1^P+M^. FUS (1.8 µM FUS supplemented with 0.2 µM FUS-GFP) droplets were formed in TAB-K^+^ ± 3 mM MgCl_2_ for 90 min prior to adding S1^P+M^ (8 or 32 µM unlabeled RNA). Images were acquired before adding RNA (“pre-RNA”) and 60 min post-RNA addition (“post-RNA”). Scale bar is 5 µm. **(J)** Representative images of permeabilized HEK 293 cells treated with RNA S1 or S1^P+M^ after sodium arsenite treatment (0.5 mM, 1 h). Cells were fixed and stained for endogenous FUS (green), TIAR (red), and DAPI (blue). Scale bar is 10 µm. **(K)** Pearson correlation of cytoplasmic FUS and cytoplasmic TIAR foci from Fig. 5J. Bars represent the mean±SEM (n=3). Large squares represent individual trial means (n=3) and small circles represent individual image means (n=75). Ordinary one-way ANOVA with Šidák’s multiple comparisons test was used to compare the indicated means (ns P>0.05, **P≤0.005, ****P<0.0001).

To dissect the contribution of each modification, we tested RNAs with only 2’-OMe (S1^M^) or phosphorothioate backbone (S1^P^) modification. At moderate concentration (8 µM) in K⁺, S1^M^ was significantly weaker, though all reversed LLPS similarly at 32 µM (**Fig. S6B-C**). In Na⁺, S1^P^ matched S1^P+M^’s potency, while S1^M^ remained weaker (**Fig. S6D-E**). Under Mg²⁺, S1 and S1^P^ activities diminished at low concentration, but S1^P^ recovered at higher levels (**Fig. S6F-G**). In contrast, S1^P+M^ and S1^M^ were unaffected by Mg²⁺ (**Fig. S6F-G**). Thermal melts confirmed that neither single modification fully stabilized S1 like S1^P+M^ (**Fig. S6H-I**). Together, these results suggest that S1^P+M^ activity is not derived from either modification alone, but rather both are needed for its enhanced activity.

In addition to stabilized rG4 structure and enhanced activity in reversing FUS LLPS, S1^P+M^ contains modifications that enhance its RNase resistance^66,67^. Therefore, we hypothesized that S1^P+M^ would have enhanced activity in cellular environments compared to S1. We utilized HEK 293 cells, where the endogenous FUS phase-separates into SG upon sodium arsenite treatment, and employed a permeabilized approach to wash away soluble protein for better visualization of phase-separated FUS^68^ and allow for the direct addition of RNA to treated cells (**Fig. 5J-K**). Without RNA, cytoplasmic FUS foci strongly colocalize with cytoplasmic TIAR foci, indicating its recruitment to stress granules (**Fig. 5J-K**; PBS). Upon RNA S1 addition, the correlation between cytoplasmic FUS and TIAR foci decreases in a concentration-dependent manner, suggesting partial disruption of FUS association with stress granules (**Fig. 5J-K**; S1). This effect is more pronounced with S1^P+M^, indicating that S1^P+M^ more effectively reduces FUS phase separation in cells compared to S1 (**Fig. 5J-K**).

## Transcriptome-wide bioinformatic analysis identifies novel RNA inhibitors of FUS LLPS and aggregation

Having identified FUS-binding rG4 as a criterion for strong RNA inhibitors and further investigated parameters governing rG4 activity, we aimed to survey the human transcriptome to identify novel rG4s capable of inhibiting FUS LLPS and aggregation. To understand how rG4 inhibitors would behave in a competitive environment, such as in the cell where other FUS-binding RNAs can interact with phase-separated FUS, and to guide our bioinformatic analysis, we reconstituted a competitive environment *in vitro.* This was done by introducing both a G4-RNA (i.e., RNA S1) and a non-G4 RNA (i.e., RNA S1_G4neg) with comparable FUS binding affinities (**Fig. S2M and P**), to pre-formed FUS droplets. We specifically selected an RNA concentration (4 µM) at which RNA S1 could partially extract FUS into the soluble phase while preserving droplet integrity (**Fig. 6A**). This setup allows us to evaluate RNA distribution between the soluble and condensed phases. When RNAs S1 or S1_G4neg were individually added to pre-formed FUS droplets, they both exhibited similar enrichment within the droplets (**Fig. 6A-B**; Single RNA). In contrast, when added together, S1_G4neg was significantly more enriched within the droplets, while RNA S1 was preferentially found in the soluble phase (**Fig. 6A-B**; Mixed RNA). This observation aligns with the distinct ability of RNA S1 to solubilize FUS and is consistent with the polyphasic linkage framework, where phase separation is weakened if the ligand (e.g., RNA) prefers to bind to sites on the scaffold (e.g., FUS) in the dilute phase vs the dense phase^69^. These results suggest that in a cellular environment containing a mixture of RNAs with high affinity for FUS, strong inhibitors are more likely to be found in the soluble FUS phase than RNAs without activity.

**Figure 6.**
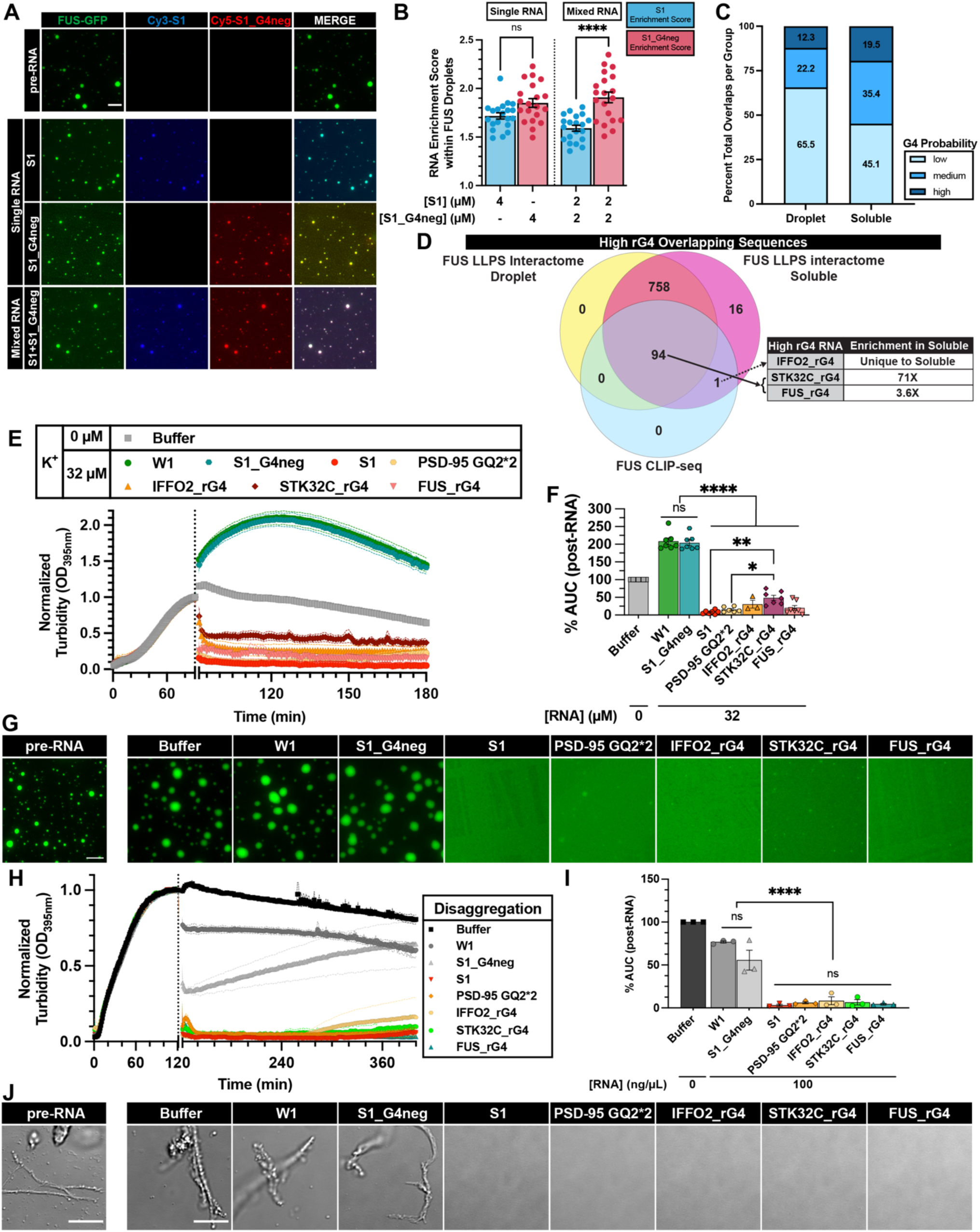
Transcriptome-wide bioinformatic analysis identifies novel RNA inhibitors of FUS LLPS and aggregation. **(B)** Representative images of FUS droplets before (“pre-RNA”, top) and after the addition of RNA S1 and/or S1_G4neg in TAB-K^+^. LLPS of FUS (1.8 µM FUS supplemented with 0.2 µM FUS-GFP) was monitored by droplet imaging before and 5 minutes after the addition of RNA. For “Single RNA”, S1 (4 µM + 75 nM Cy3-S1) or S1_G4neg (4 µM + 75 nM Cy5-S1_G4neg) were individually added to pre-formed FUS droplets. For “Mixed RNA”, S1 (2 µM + 75 nM Cy3-S1) and S1_G4neg (2 µM + 75 nM Cy5-S1_G4neg) were added to pre-formed FUS droplets. Scale bar is 5 µm. **(C)** Average RNA enrichment score within FUS droplets was quantified for the post-RNA images from Fig. 6A. Bars represent mean±SEM and data points represent individual image averages (n=20) across two independent trials. Ordinary one-way ANOVA with Šidák’s multiple comparisons test was used to compare the means of different RNA (ns P>0.05, ****P<0.0001). **(D)** Intersection of FUS phase separation-dependent interactome (Droplet or Soluble)^39^ and the transcriptome-wide rG4 database^43^. Overlaps with high probability of forming rG4 are in dark blue, those with medium probability are in blue, and those with low probability are in light blue. **(E)** Venn diagram of the high-probability rG4 sequences that overlap between FUS Droplet interactome, FUS Soluble interactome, and/or FUS CLIP-seq^48^ interactome. Indicated RNA sequences were selected from this analysis and used in Fig. 6E-H and Fig. S7F-I. **(F)** Turbidity measurement of FUS LLPS reversal with RNA in TAB-K^+^. LLPS of FUS (2 µM) was initiated, and monitored by turbidity at 395 nm. After 90 min, data collection was paused to add RNA (32 µM). Data represent mean±SEM (n=3-7). **(G)** AUC was calculated from turbidity curves in Fig. 6E after RNA was added. Data represent mean±SEM (n=3-7). Ordinary one-way ANOVA with Tukey’s multiple comparisons test was used to compare the means between RNAs (ns P>0.05, *P≤0.05, **P≤0.005, ****P<0.0001). **(H)** Representative images of FUS droplets before (“pre-RNA”) and after the addition of indicated RNAs. LLPS of FUS (1.8 µM FUS supplemented with 0.2 µM FUS-GFP) was monitored by droplet imaging before and 60 minutes after the addition of RNA (32 µM unlabeled RNA). Scale bar is 5 µm. **(I)** Turbidity measurement of FUS disaggregation with indicated rG4 RNAs and non-rG4 RNA controls. FUS (5 µM) aggregation was initiated by adding TEV protease to GST-TEV-FUS in Assembly Buffer and monitored by turbidity at 395 nm. After 120 min, data collection was paused to add RNA (100 ng/µL). Data represent mean±SEM (n=3). **(J)** AUC was calculated from turbidity curves in Fig. 6H after RNA was added. Data represent mean±SEM (n=3). Ordinary one-way ANOVA with Tukey’s multiple comparisons test was used to compare the means between the RNA treatments. (ns P>0.05, ****P<0.0001). **(K)** Representative differential interference contrast (DIC) images of FUS aggregates before and after the addition of RNA. Pre-formed FUS aggregates were assembled as in (H), and images were acquired immediately before (“pre-RNA”) or 100 min after addition of buffer or RNA (100 ng/µL). Scale bar is 10 µm.

Guided by this observation, we performed a bioinformatic analysis to identify FUS-bound rG4s enriched in the soluble FUS phase. We queried the intersection of three datasets: transcriptome-wide rG4 database (GSE77282)^43^, FUS CLIP-seq dataset (GSE40651, sample GSM998875)^48^, and the phase separation-dependent FUS interactome (E-MTAB-8456)^39^. The rG4 database is particularly well-suited for this analysis because all entries consist of 30-nucleotide RNAs, a length comparable to our known RNA inhibitors, and likely represents an optimal size for RNA function (see discussion below). We first compared the rG4 database and the FUS CLIP database and found 95 FUS-binding sequences overlapping with the high-probability rG4 database (**Fig. S7A**). Notably, this overlap is significantly higher than that between the rG4 database and the TAF15 CLIP database (26 high-probability rG4 overlaps) (GSE43294)^47^, indicating a preferential binding of high-probability rG4s to FUS compared to its FET family member TAF15 (**Fig. S7A**). Further comparison of the rG4 database with the FUS phase-dependent interactome showed an enrichment of high-probability rG4s in the soluble FUS phase (**Fig. 6C**), consistent with the result of our competitive assay (**Fig. 6B**). Moreover, of the 870 high-probability rG4-forming sequences in the rG4 database^43^, 869 were found in the soluble FUS interactome^39^, indicating FUS has a strong recognition of rG4. To confirm this enrichment was not due to the promiscuous RNA-binding ability of FUS, we performed two-way permutation tests with 100 sets of random transcripts matched for number and length of the rG4 database and FUS phase-dependent interactome (**Fig. S7B-E**), validating the statistical significance of this enrichment and further supporting the selectivity of FUS to rG4.

We then cross-referenced the 95 FUS-binding, high-probability rG4s with the 869 high-probability rG4s from the FUS soluble-phase interactome to identify the FUS-binding rG4s that are most enriched in the soluble phase, which we reason would be strong inhibitors of FUS LLPS (**Fig. 6D)**. One sequence (i.e., IFFO2_rG4) was found exclusively in the soluble FUS interactome and was thus selected for further testing. Among the remaining 94 sequences present in both phases, STK32C_rG4 showed the greatest enrichment in the soluble phase (see Methods) and was also chosen (**Fig. 6D**). Interestingly, FUS was also found to bind an rG4 sequence in its own mRNA that is enriched in the soluble phase, and this sequence (FUS_rG4) was included as well. After validating that this panel of RNAs adopts rG4 structure by CD (**Fig. S7F**), we assessed their ability to reverse FUS LLPS and aggregation. Remarkably, all three FUS-binding rG4s have activity similar to strong inhibitors RNA S1 and length-matched PSD-95 GQ2*2, as they can strongly reverse FUS LLPS (**Figs. 6E-G**) and aggregation (**Figs. 6H-J, S7G-H**). Together, these results establish that short rG4s maintain FUS solubility by engaging the diffuse phase, providing a mechanistic basis for identifying novel potent RNA inhibitors of FUS LLPS and aggregation.

### DISCUSSION

FUS phase separation is essential for its physiological roles in RNA metabolism and cellular stress response. However, dysregulation of this process and subsequent protein aggregation are strongly linked to neurodegenerative diseases, including amyotrophic lateral sclerosis (ALS) and frontotemporal dementia (FTD). Identifying molecular features that govern FUS LLPS is therefore critical for understanding disease mechanisms and developing targeted therapeutic strategies. While RNA is known to regulate FUS condensation^3,27,30^, the sequence and structural rules that distinguish inhibitory from promoting RNAs remain poorly defined. Here, we demonstrate that RNA G-quadruplexes (rG4s) serve as key structural motifs that enable potent and tunable inhibition of FUS phase separation. Modulating rG4 length or stability precisely alters RNA activity, establishing a direct structure-function relationship. Although rG4s interact with both soluble and condensed FUS, they preferentially engage the soluble pool, likely shifting the phase equilibrium toward dispersion^69^. Guided by these mechanistic insights, we identified a new class of short, endogenous rG4 RNAs that potently reverse FUS LLPS and aggregation.

## Mechanisms of rG4-mediated reversal of FUS LLPS: rG4 concentration, stability, and length

One key distinction between strong and weak RNA inhibitors is that strong inhibitors exhibit concentration-dependent activities against FUS LLPS, whereas weak inhibitors have limited activity regardless of concentration. Additionally, strong RNA inhibitors can be highly sensitive to salt conditions, with their activity enhanced in rG4-stabilizing buffers (K^+^ >> Na^+^) and diminished in rG4-destabilizing buffers (Mg^2+^). This sensitivity is due to salt-induced structural changes in strong inhibitors, reinforcing that rG4 is a critical structural determinant of RNA activity.

Strong RNA inhibitors, such as S1, exhibit higher FUS-binding affinity than weak inhibitors, like W1^30^. In this study, we further demonstrate that binding affinity alone does not dictate RNA activity, but rather RNA structure is an equally important factor. For instance, RNA S1 and its G4-deficient counterpart (S1_G4neg) display distinct inhibitory effects, although they bind FUS with similar affinity, indicating possibly different binding patterns^64,70,71^. Notably, while S1_G4neg preferentially binds to phase-separated FUS, S1 favors soluble FUS, potentially shifting the equilibrium toward the dispersed phase and promoting droplet dissolution^69^.

Another key factor influencing RNA function is the repeat length of the rG4 element. We demonstrated that RNAs with one or two repeats of rG4-forming RNA S1 effectively inhibit FUS LLPS. However, increasing the repeat number to ten shifts its function from inhibition to promotion of phase separation (**Figs. 4, S5**). This finding aligns with previous studies showing that rG4 can promote the aggregation of neurodegenerative disease proteins such as TDP-43^72^, α-synuclein^73^ and tau^74^. Our results highlight a protective role of short rG4 in preventing FUS assembly and suggest that repeat length is a crucial factor in determining whether an RNA functions as an inhibitor or nucleator of FUS aggregation. Future studies should aim to define the precise critical length for different proteins at which this transition occurs.

Our findings further suggest that rG4 stability plays a key role in mitigating FUS phase separation. RNAs with chemical modifications conferring stronger rG4-forming potential, such as S1^P+M^, exhibit strong reversal activity and greater resistance to salt-induced activity changes. Previous studies have reported that FUS binding can both destabilize^58,64^ and stabilize rG4 structures^75^. On the other hand, rG4 could also reshape the protein folding energy landscapes^76^, potentially further contributing to the activities of rG4 in regulating FUS LLPS, and warrants future investigation.

## Considerations when developing short RNAs as therapeutics to mitigate aberrant FUS phase transition

Our previous study has demonstrated that short RNA oligonucleotides, such as RNA S1, can reverse neuronal toxicity by mitigating aberrant FUS phase transition^30^. Thus, strong RNA inhibitors like RNA S1 have the potential to be developed as therapeutics for ALS/FTD that are characterized by FUS aggregation. The mechanistic insight provided by the current study is critical for further development of these therapeutic RNA oligonucleotides. We identified parallel rG4 as a defining feature of strong RNA inhibitors of FUS LLPS. Another critical consideration is the concentration-dependent activity of strong RNA inhibitors, which underscores the need for thorough optimization of dosage when designing RNA therapeutics.

Furthermore, our findings suggest that therapeutic RNA design must account for RNA structural stability under varying ionic conditions. The structure and activity of rG4-containing RNAs, such as RNA S1 and S2, are highly sensitive to buffer conditions, with their ability to inhibit FUS LLPS completely abolished in magnesium-containing buffers. Moreover, recent studies have demonstrated that calcium can significantly alter the phase behavior of rG4-containing RNAs^59,73^. These findings are particularly important because both calcium and magnesium are present in neurons, potentially limiting the efficacy of therapeutic RNAs in the cellular environment. Importantly, our study provides insight into strategies for overcoming this structural sensitivity. We found that modifications such as phosphorothioate backbones and 2’-OMe groups significantly enhance rG4 stability, rendering RNA S1 resistant to environmental changes. Interestingly, these modifications also improve the RNA’s resistance to RNase. Thus, they may improve the robustness of rG4-based therapeutic RNAs in physiological conditions. Indeed, we showed that RNA S1 with these modifications (RNA S1^P+M^) has enhanced activity against FUS phase separation in cells (**Fig. 5**). Additionally, structural features such as loop length, tetrad number, and intramolecular rG4 formation further contribute to rG4 stability ^77,78^. Future studies will explore these factors to develop robust rG4-containing therapeutic RNAs.

Our study also provides valuable insights into the optimal length of therapeutic RNAs. For RNA S1, a single repeat (25 nucleotides) appears to be optimal, as doubling the length, despite providing structural stability, does not enhance activity when compared at the same mass concentration (**Fig. 4**). Further increasing the repeat length to 10 repeats (252 nucleotides, RNA S1*10) even converts the RNA into a nucleator of FUS assembly. This observation is consistent with previous findings involving polyU RNAs, which demonstrated that polyU sequences of 30 nucleotides or less form complexes with a single unit of FUS, whereas longer polyU RNAs form complexes with two or three FUS units^14^. Collectively, these results suggest that short, strong rG4-forming RNAs of approximately 25-30 nucleotides, combined with appropriate chemical modifications to stabilize the rG4 structure, represent promising candidates for therapeutic development.

## Identifying novel rG4 regulators of FUS phase separation from the human transcriptome

Using FUS-binding rG4s as a selection criterion and focusing on short, high-probability rG4s enriched in the soluble phase of the phase separation-dependent FUS interactome, our bioinformatic analysis identified novel rG4 regulators of FUS phase separation from the human transcriptome. The rG4 database was particularly well-suited for this analysis, as all entries are 30-nucleotide RNAs, which is within the optimal length for RNA function. Notably, the FUS phase separation-dependent interactome contains the majority of high-probability rG4s in the database, indicating strong recognition by FUS. FUS engages endogenous rG4 RNAs in cells, and these interactions serve diverse biological roles. For example, FUS interacts with TERRA RNA to regulate telomere length^75^ and interacts with G_4_C_2_ RNA to inhibit its RAN translation^64^. Recently, it was shown that expression of ALS-associated mutation FUS-P525L reduces the recruitment of rG4 into stress granules^46^. Despite these insights, the impact of rG4-mediated phase regulation on FUS function remains poorly understood. Future studies are needed to elucidate the functional interplay between rG4 activity and FUS phase behavior in cells, and our study has identified important targets (**Fig. 6D-J**) for these future studies.

In totality, our findings position rG4-containing RNAs as a new class of chemically tunable regulators of FUS phase behavior. By linking RNA secondary structure to a concentration- and stability-dependent function in LLPS regulation, this work establishes a design framework for engineering short RNAs that either suppress or promote biomolecular condensation. The discovery that rG4 length acts as a molecular switch, converting RNAs from inhibitors to nucleators, introduces a powerful structural axis for programming phase transitions. Moreover, chemical modifications that enhance rG4 stability not only boost functional potency but also confer robustness in complex cellular environments. These insights offer a foundation for developing rG4-based therapeutics for FUS-driven neurodegeneration and suggest broader applications for modulating other phase-separating systems. More broadly, this work demonstrates that structured RNAs can be rationally designed to reshape protein phase landscapes, opening new frontiers at the interface of RNA chemistry, condensate biology, and disease intervention.

## ACKNOWLEDGEMENTS

L.G. is supported by Dr. Ralph and Marian Falk Medical Research Trust, Frick Foundation for ALS Research, the National Institute of General Medical Sciences grant R35GM138109, and the National Institute of Neurological Disorders and Stroke grant R01NS121143. J.S. is supported by grants from The Packard Center for ALS Research at Johns Hopkins, Target ALS, The Association for Frontotemporal Degeneration, the Amyotrophic Lateral Sclerosis Association, the Office of the Assistant Secretary of Defense for Health Affairs through the Amyotrophic Lateral Sclerosis Research Program (W81XWH-20-1-0242 and W81XWH-17-1-0237), and NIH grant R01GM099836. Z.S and H.W are supported by the National Institute of General Medical Sciences grant R35GM147027. J.A.D. and K.B. are supported by T32GM144302. A.R.H. is supported by NIH grants RF1NS114128, R01NS114128, and the philanthropic support of the Farber Family Foundation to the Jefferson Weinberg ALS Center.

## AUTHOR CONTRIBUTIONS

J.L.C., M.H., L.R.G., H.W., K.B., and J.A.D. performed the experimental studies and carried out the analysis. E.W. performed the bioinformatic analysis. J.L.C. and L.G. wrote the original draft of the manuscript with input and edits from all authors. Z.S., J.S., S.M., A.R.H., and L.G. supervised the work.

## SUPPLEMENTARY DATA

Supplementary Data are available online.

## DATA AVAILABILITY

All data presented in this study are available upon request to the corresponding authors.

