## Supplementary Figures and Tables for "RNA G-Quadruplexes Function as a Tunable Switch of FUS Phase Separation"

### SUPPLEMENTARY FIGURES and LEGENDS

Supplementary Figure 1

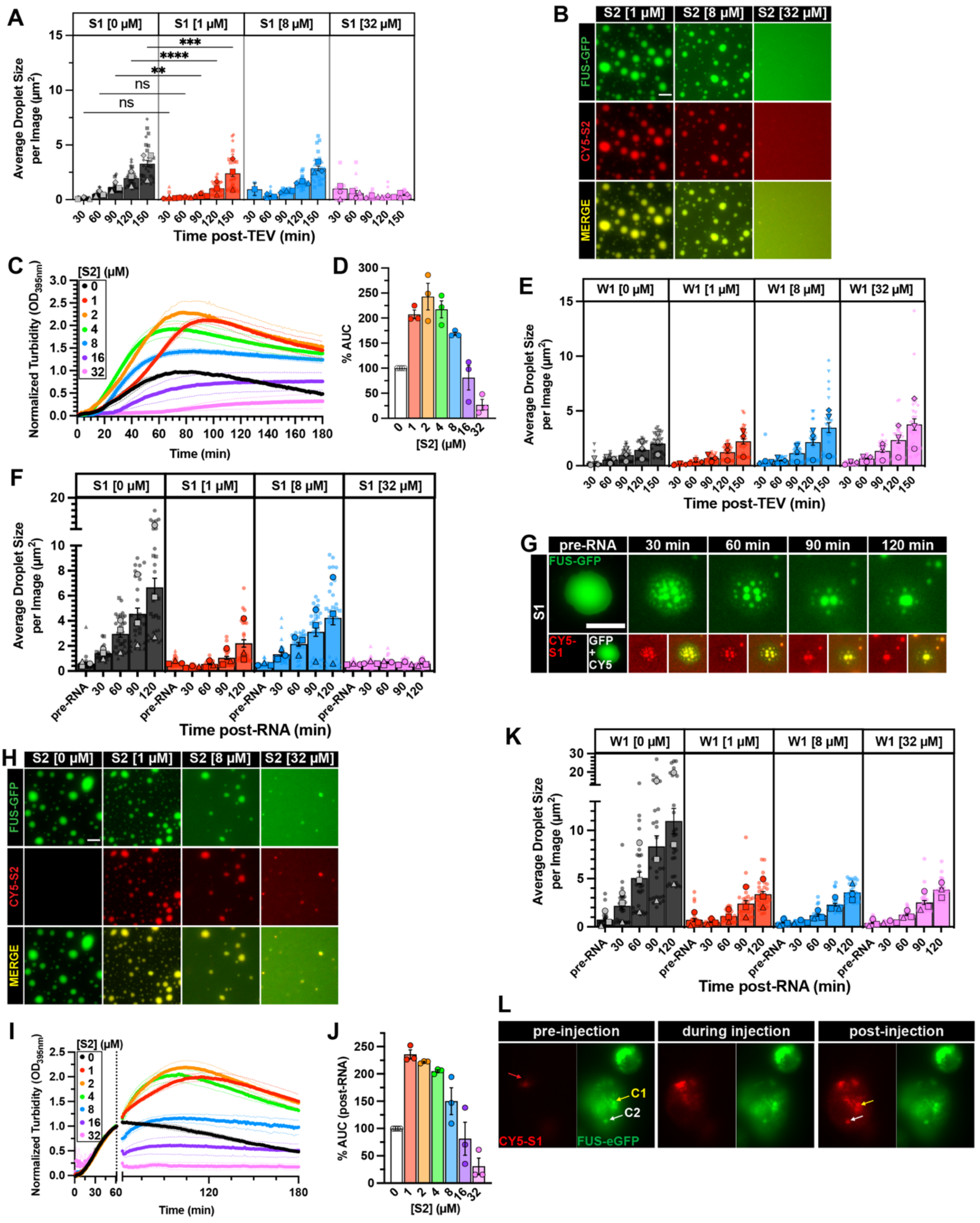

**Supplementary Figure 1. Strong RNA inhibitors prevent and reverse FUS phase separation in a concentration-dependent manner.**

(**A,E,F,K**) Average droplet size was quantified from images collected in Figures 1B, D (**A**) or Figure 1G (**E**), or from reversal images collected in Fig. 1K (**F**), or Fig. 1O (**K**). Individual points represent image means (n=30), large symbols represent trial means (n=3), and bars represent the mean±SEM of three independent experiments. For Fig. S1A, ordinary one-way ANOVA with Šidák's multiple comparisons test was used to compare the means between 0 µM S1 and 1 µM S1 droplets at each time point (ns P>0.05, **P≤0.005, ***P≤0.0005, ****P<0.0001).

(**B**) Representative images of FUS droplets formed in the presence of RNA S2. LLPS of FUS (1.8 µM FUS supplemented with 0.2 µM FUS-GFP) with S2 (1, 8, or 32 µM unlabeled S2 supplemented with 75 nM Cy5-S2) was monitored by droplet imaging every 30 minutes for a duration of 150 minutes. The 120-minute timepoint images were selected to represent RNA effect on FUS LLPS. Scale bar is 5 µm.

(**C**) Turbidity measurement of FUS LLPS inhibition by RNA S2. MBP-FUS (2 µM) was incubated in the presence of varying concentrations of S2 prior to initiating FUS LLPS reaction with TEV protease. Turbidity at 395 nm was monitored at 25°C over 180 min. Data represent mean±SEM (n=3-4). The 0 µM RNA control plotted in Fig. 1E, 1I, and S1C are the same because these experiments were conducted simultaneously.

(**D**) Area under the curve (AUC) was calculated from turbidity curves in Fig. S1C. Data represent mean±SEM (n=3-4).

(**G**) Zoomed representative images of the same FUS droplets before and after the addition of RNA S1 to illustrate strong RNA "breaking apart" a large and highly-enriched FUS droplet into numerous smaller droplets. FUS (1.8 µM FUS supplemented with 0.2 µM FUS-GFP) droplets were formed for 60 minutes prior to adding S1 (32 µM unlabeled S1 supplemented with 75 nM Cy5-S1). Scale bar is 5 µm.

(**H**) Representative images of FUS droplet 60 minutes after addition of RNA S2. FUS (1.8 µM FUS supplemented with 0.2 µM FUS-GFP) droplets were formed for 60 minutes prior to adding S2 (1, 8, or 32 µM unlabeled S2 supplemented with 75 nM Cy5-S2). Droplets were monitored by imaging immediately before RNA, then every 30 minutes for a duration of 120 minutes after RNA was added. The 60-minute post-RNA timepoint images were selected to represent RNA's effect on FUS LLPS. Scale bar is 5 µm.

(**I**) Turbidity measurement of FUS LLPS reversal with RNA S2. FUS (2 µM) LLPS was initiated, and monitored by turbidity at 395 nm. After 60 min, data collection was paused to add S2, then resumed to monitor turbidity changes for an additional 120 min. Data represent mean±SEM (n=3-4).

(**J**) AUC was calculated from turbidity curves in Fig. S1I after RNA was added. Data represent mean±SEM (n=3-4).

(**L**) Microinjection of Cy5-RNA S1 (0.5 µM) into HEK 293T cells expressing FUS-WT-eGFP. The red arrow marks the site of RNA injection. White and yellow arrows mark Cy5-S1 localization to cytoplasmic FUS condensates ("C1" or "C2") post-injection.

Supplementary Figure 2

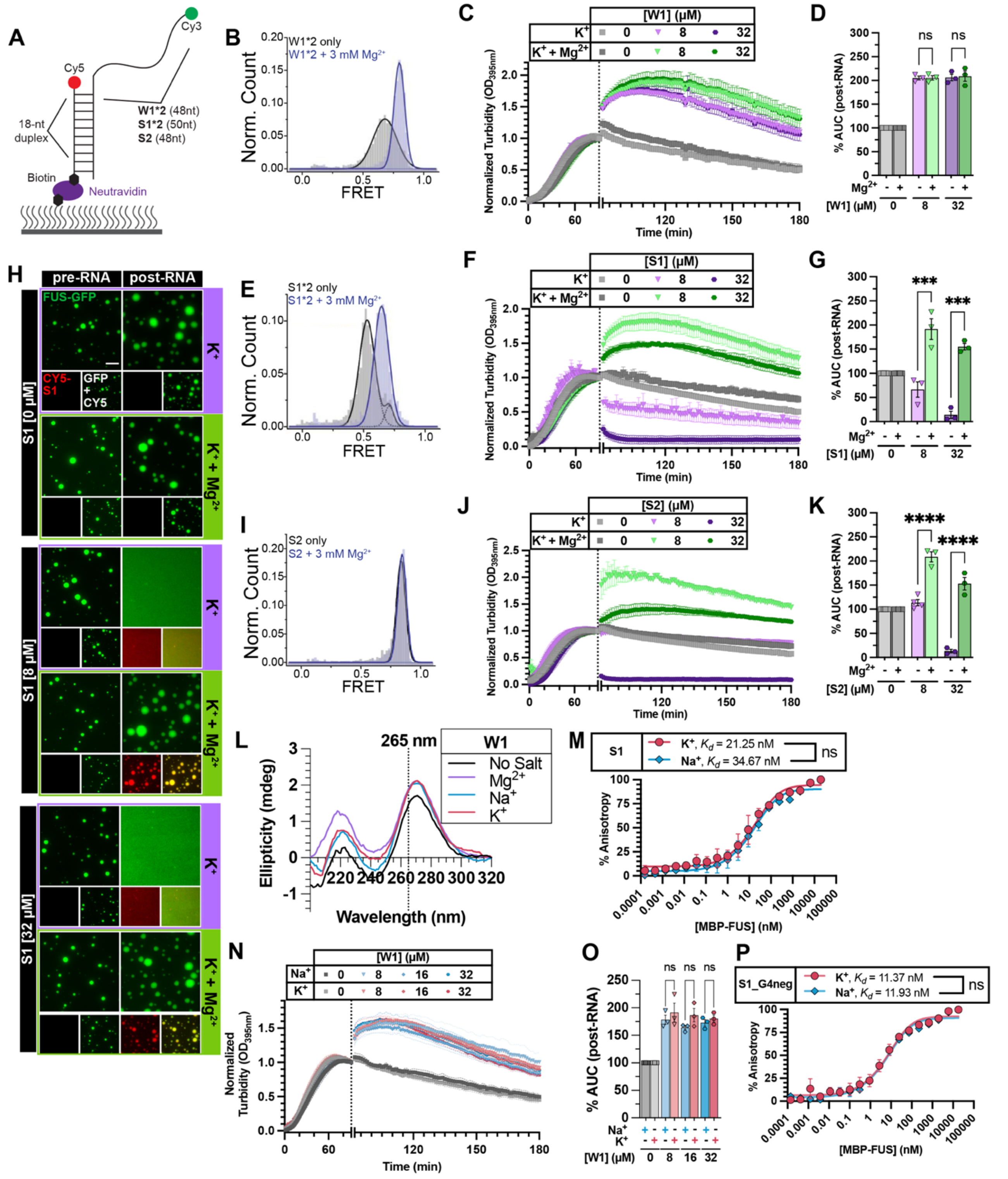

**Supplementary Figure 2. rG4 is the specific structural determinant for RNA S1 activity in mitigating FUS LLPS.**

(**A**) smFRET experimental diagram. RNA W1 (24nt) and S1 (25nt) were doubled (W1*2 or S1*2) to achieve optimal length for smFRET experiments.

(**B, E, I**) FRET efficiency histogram determined by smFRET experiment for RNA W1*2 (**B**), RNA S1*2 (**E**), or RNA S2 (**I**) in the presence of 100 mM KCl (black trace) or 100 mM KCl supplemented with 3 mM $MgCl_2$ (blue trace).

(**C, F, J**) Turbidity measurement of FUS LLPS reversal with RNA W1 (**C**), S1 (**F**), or S2 (**J**) in TAB-$K^+$ ± $Mg^{2+}$. FUS (2 μM) LLPS was initiated, and monitored by turbidity at 395 nm. After 90 min, data collection was paused to add RNA diluted in 10 mM Tris-HCl pH 7.4 supplemented with 50 mM KCl (purple symbols) or 50 mM KCl + 3 mM $MgCl_2$ (green symbols). Data represent mean±SEM (n=3). The 0 μM $K^+$ control traces are the same for the following because these experiments were conducted simultaneously: Figs. 2D & S2F or Figs. S2J & S3C.

(**D, G, K**) AUC was calculated from turbidity curves in Fig. S2C (**D**), Fig. S2F (**G**), or Fig. S2J (**K**) after RNA was added. Data represent mean±SEM (n=3). Ordinary one-way ANOVA with Šidák's multiple comparisons test was used to compare the means (ns P>0.05, ***P≤0.0005, ****P<0.0001).

(**H**) Representative images of FUS droplets in TAB-$K^+$ ± 3 mM $MgCl_2$ before and after the addition of RNA S1. FUS (1.8 μM FUS supplemented with 0.2 μM FUS-GFP) droplets were formed in TAB-$K^+$ ± 3 mM $MgCl_2$ for 90 minutes prior to adding S1 (8 or 32 μM unlabeled RNA supplemented with 75 nM Cy5-RNA). Images were acquired before adding RNA ("pre-RNA") and 60 min post-RNA addition ("post-RNA"). Scale bar is 5 μm.

(**L**) Representative CD spectra of RNA W1 collected at 25°C in indicated buffer condition.

(**M, P**) Anisotropy measurement of fluorescein (fl)-RNA S1 (**M**) or fluorescein-RNA S1_G4neg (**P**) with MBP-FUS. For each trial, fl-RNA (8 nM) was folded then added to increasing concentrations of MBP-FUS in TAB-$Na^+$ (blue diamonds) or TAB-$K^+$ (red circles). Fluorescence anisotropy was normalized, and the mean±SEM of three independent trials was plotted. Solid lines represent the fitted curve to obtain $K_d$. Extra sum-of-squares F Test was used to compare $K_d$ in TAB-$Na^+$ and TAB-$K^+$ (ns P>0.05).

(**N**) Turbidity measurement of FUS LLPS reversal with W1 in TAB-$Na^+$ (blue traces) or TAB-$K^+$ (red traces). LLPS of FUS (2 μM) was initiated, and monitored by turbidity at 395 nm. After 90 min, data collection was paused to add W1. Increasing concentration of RNA is indicated by a more saturated color. Data represent mean±SEM (n=3).

(**O**) AUC was calculated from turbidity curves in Fig. S2N after RNA was added. Data represent mean±SEM (n=3). Ordinary one-way ANOVA with Šidák's multiple comparisons test was used to compare the means between TAB-$Na^+$ and TAB-$K^+$ at each RNA concentration (ns P>0.05).

Supplementary Figure 3

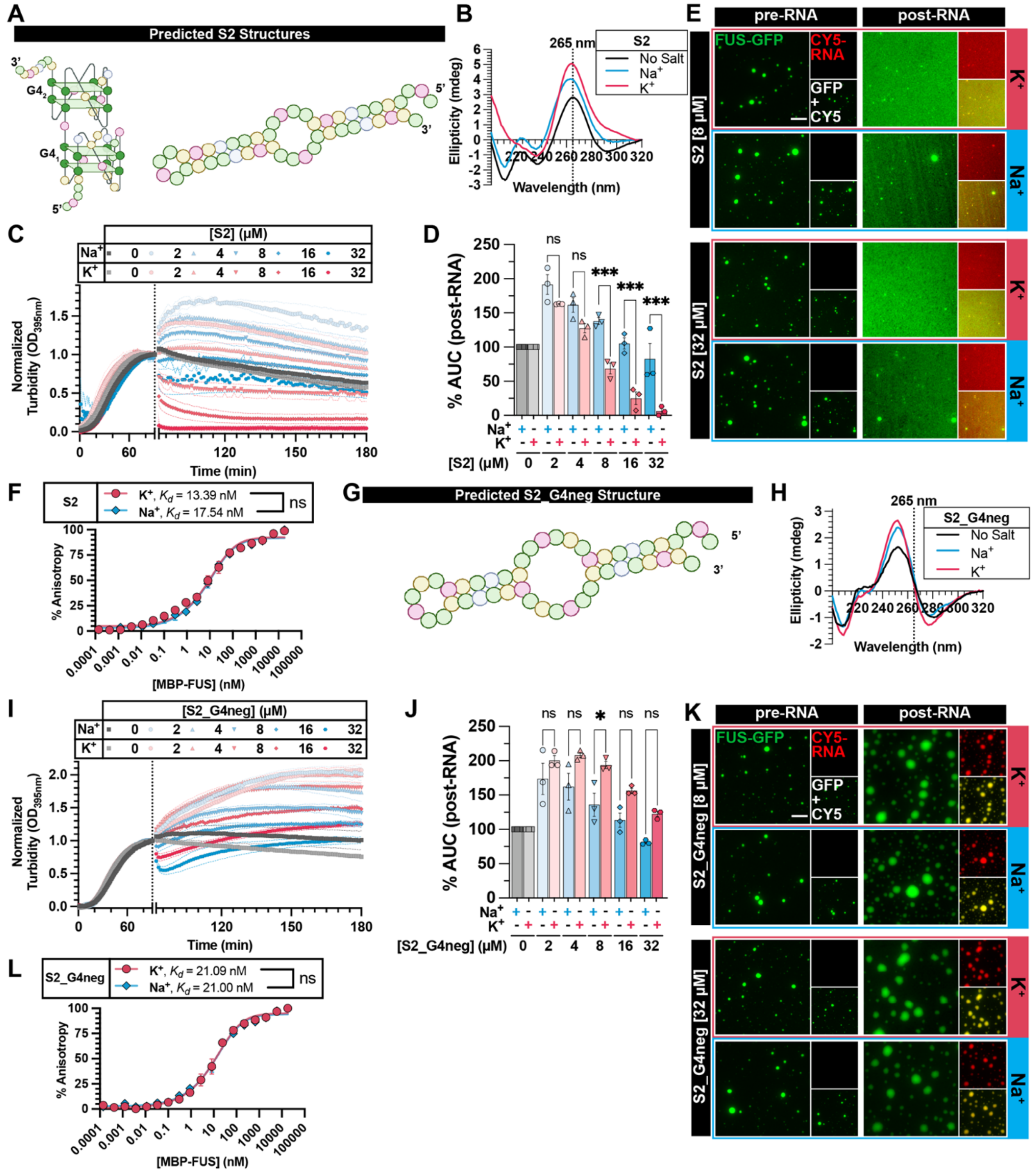

**Supplementary Figure 3. rG4, not hairpin, structure also determines RNA S2 activity in mitigating FUS LLPS.**

(**A, G**) Schematic representations of predicted rG4 and/or stem-loop structures formed by RNA S2 (**A**) or S2_G4neg (**G**). The rG4 structure was predicted using QGRS Mapper[1] , and the canonical secondary structure was predicted using RNAfold WebServer[2]. RNA structures were created with Biorender.com.

(**B, H**) CD spectra of RNA S2 (**B**) or S2_G4neg (**H**) collected at 25°C in indicated buffer condition.

(**C, I**) Turbidity measurement of FUS LLPS reversal with RNA S2 (**C**) or S2_G4neg (**I**) in TAB-$Na^+$ (blue traces) or TAB-$K^+$ (red traces). LLPS of FUS (2 μM) was initiated, and monitored by turbidity at 395 nm. After 90 min, data collection was paused to add RNA. Increasing concentration of RNA is indicated by a more saturated color. Data represent mean±SEM (n=3). The 0 μM RNA control in TAB-$K^+$ plotted in Fig. S3C and Fig. S2J are the same because these experiments were conducted simultaneously.

(**D, J**) AUC was calculated from turbidity curves in Fig. S3C (**D**) or Fig. S3I (**J**) after RNA was added. Data represent mean±SEM (n=3). Ordinary one-way ANOVA with Šidák's multiple comparisons test was used to compare the means between TAB-$Na^+$ and TAB-$K^+$ at each RNA concentration (ns $P>0.05$, *$P\leq0.05$, ***$P\leq0.0005$).

(**E, K**) Representative images of FUS droplets before and after the addition of RNA S2 (**E**) or S2_G4neg (**K**). LLPS of FUS (1.8 μM FUS supplemented with 0.2 μM FUS-GFP) was monitored by droplet imaging before and 60 minutes after the addition of RNA (8 or 32 μM unlabeled RNA supplemented with 75 nM Cy5-RNA). Scale bar is 5 μm.

(**F, L**) Anisotropy measurement of fluorescein-RNA S2 (**F**) or fluorescein-RNA S2_G4neg (**L**) with MBP-FUS. For each trial, fl-RNA (8 nM) was folded then added to increasing concentrations of MBP-FUS in TAB-$Na^+$ (blue diamonds) or TAB-$K^+$ (red circles). Fluorescence anisotropy was normalized, and the mean±SEM of three independent trials was plotted. Solid lines represent the fitted curve to obtain $K_d$. Extra sum-of-squares F Test was used to compare $K_d$ in TAB-$Na^+$ and TAB-$K^+$ (ns $P>0.05$).

Supplementary Figure 4

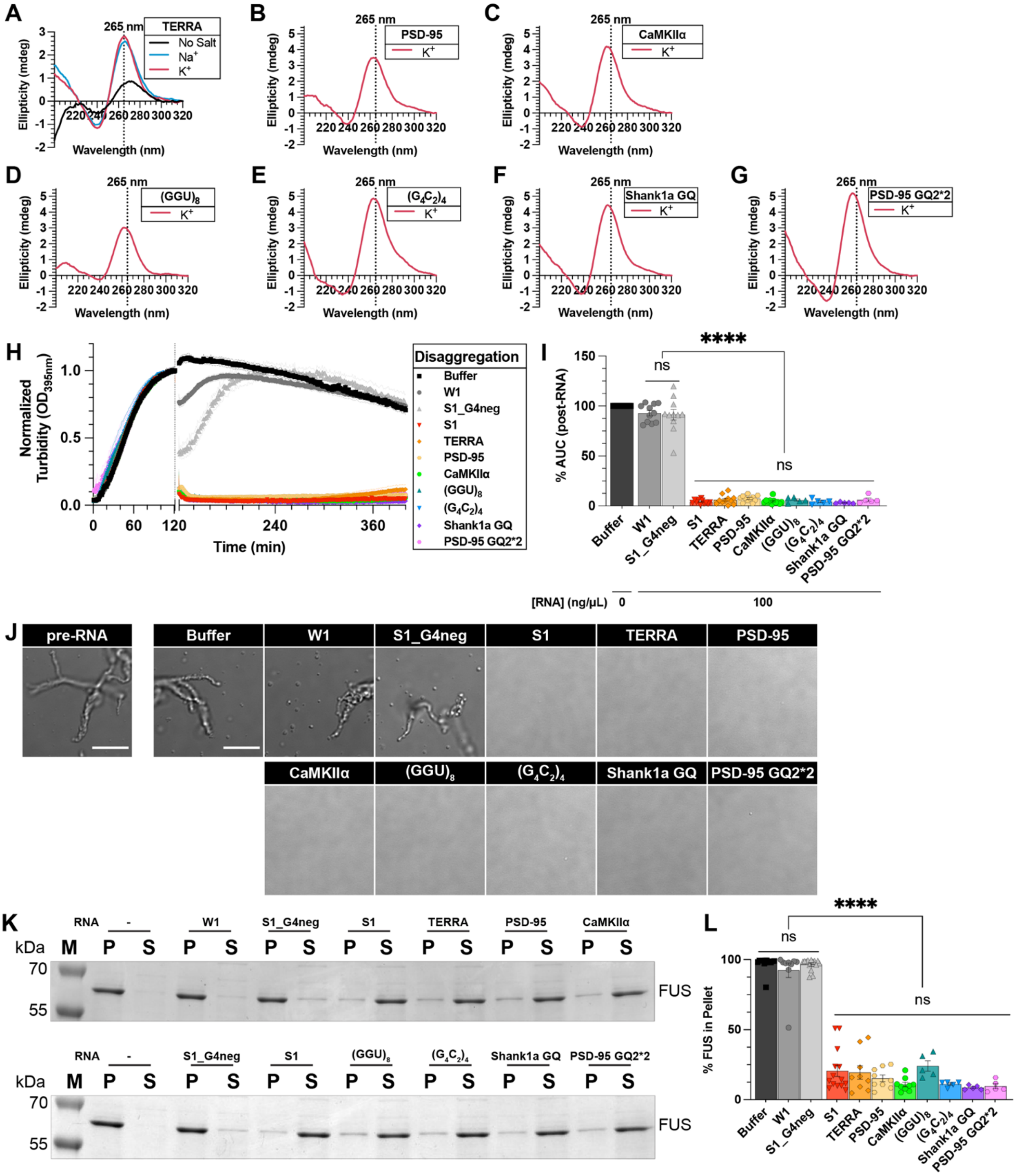

**Supplementary Figure 4. FUS-binding rG4 is a defining criterion for identifying strong RNA inhibitors of FUS LLPS and aggregation.**

(**A-G**) Representative CD spectra of RNA TERRA (**A**), PSD-95 (**B**), CaMKIIα (**C**), $(GGU)_8$ (**D**), $(G_4C_2)_4$ (**E**), Shank1a GQ (**F**), or PSD-95 GQ2*2 (**G**) collected at 25°C.

(**H**) Turbidity measurement of FUS disaggregation with indicated rG4 RNAs and non-rG4 RNA controls. FUS (5 μM) aggregation was initiated by adding TEV protease in GST-TEV-FUS in

Assembly Buffer and monitored by turbidity at 395 nm. After 120 min, data collection was paused to add RNA (100 ng/μL). Data represent mean±SEM (n=5-11).

(**I**) AUC was calculated from turbidity curves in Fig. S4H after RNA was added. Data represent mean±SEM (n=5-11). Ordinary one-way ANOVA with Tukey's multiple comparisons test was used to compare the means between the RNA treatments. (ns P>0.05, ****P<0.0001).

(**J**) Representative differential interference contrast (DIC) images of FUS aggregates before and after the addition of RNA. Pre-formed FUS aggregates were assembled as in Fig. S4H and images were acquired before ("pre-RNA") or 90 min after addition of buffer or RNA (100 ng/μL). Scale bar is 10 μm.

(**K**) Representative sedimentation analysis gels. FUS (5 μM) aggregation was initiated as in Fig. S4H. After 120 min, RNAs (100 ng/μL) were added and incubated at room temperature. Samples were collected 120 min after the addition of buffer or RNA and analyzed by sedimentation followed by SDS-PAGE.

(**L**) Quantification of SDS-PAGE gel images from Fig. S4K. Data represent mean±SEM (n=5-9). Ordinary one-way ANOVA with Tukey's multiple comparisons test was used to compare the means between the conditions. (ns P>0.05, ****P<0.0001).

Supplementary Figure 5

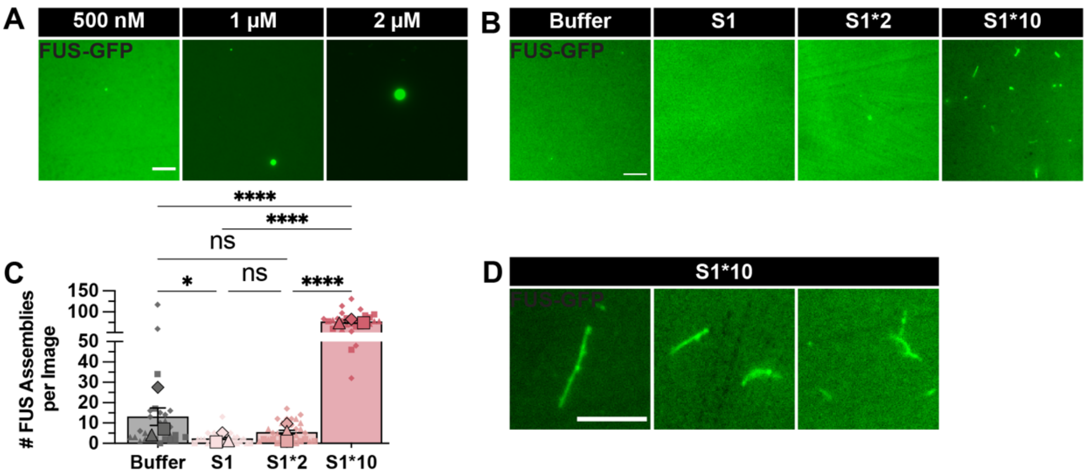

**Supplementary Figure 5. Increasing rG4 repeat numbers in RNA S1 switches it from inhibiting to nucleating FUS assembly.**

(**A**) Representative images of FUS droplets formed by varying concentrations of FUS-GFP in TAB-$K^+$. LLPS of FUS-GFP (500 nM, 1 µM, or 2 µM) in TAB-$K^+$ was initiated by adding TEV protease to cleave MBP tag from MBP-TEV-FUS-GFP. Images were acquired 120 minutes after TEV addition. Scale bar is 5 µm.

(**B**) Representative images of FUS assembly formed with 200 nM of S1, S1*2, or S1*10 in TAB-$K^+$. LLPS of FUS-GFP (500 nM) in TAB-$K^+$ was initiated as in (A) in the presence of indicated RNAs. For each trial, images were acquired 120 minutes after TEV addition (n=3). Scale bar is 5 µm.

(**C**) The average number of FUS assemblies per image was quantified from images collected in Fig. S5B and Fig. 4J. Bars represent mean±SEM of three independent experiments, large symbols represent trial means (n=3), and small datapoints represent individual image values (n=30). The 0 µM RNA control and S1*10 (200 nM) data plotted in Fig. S5C and Fig. 4K are the same because these experiments were conducted simultaneously. Ordinary one-way ANOVA with Tukey's multiple comparisons test was used to compare the means between RNAs. (ns P>0.05, ****P<0.0001).

(**D**) Representative images of FUS assemblies formed in the presence of S1*10. LLPS of FUS-GFP (500 nM) in TAB-$K^+$ was initiated as in (A) in the presence of S1*10 (200 nM). Images were acquired 120 minutes after TEV addition. Scale bar is 5 µm.

Supplementary Figure 6

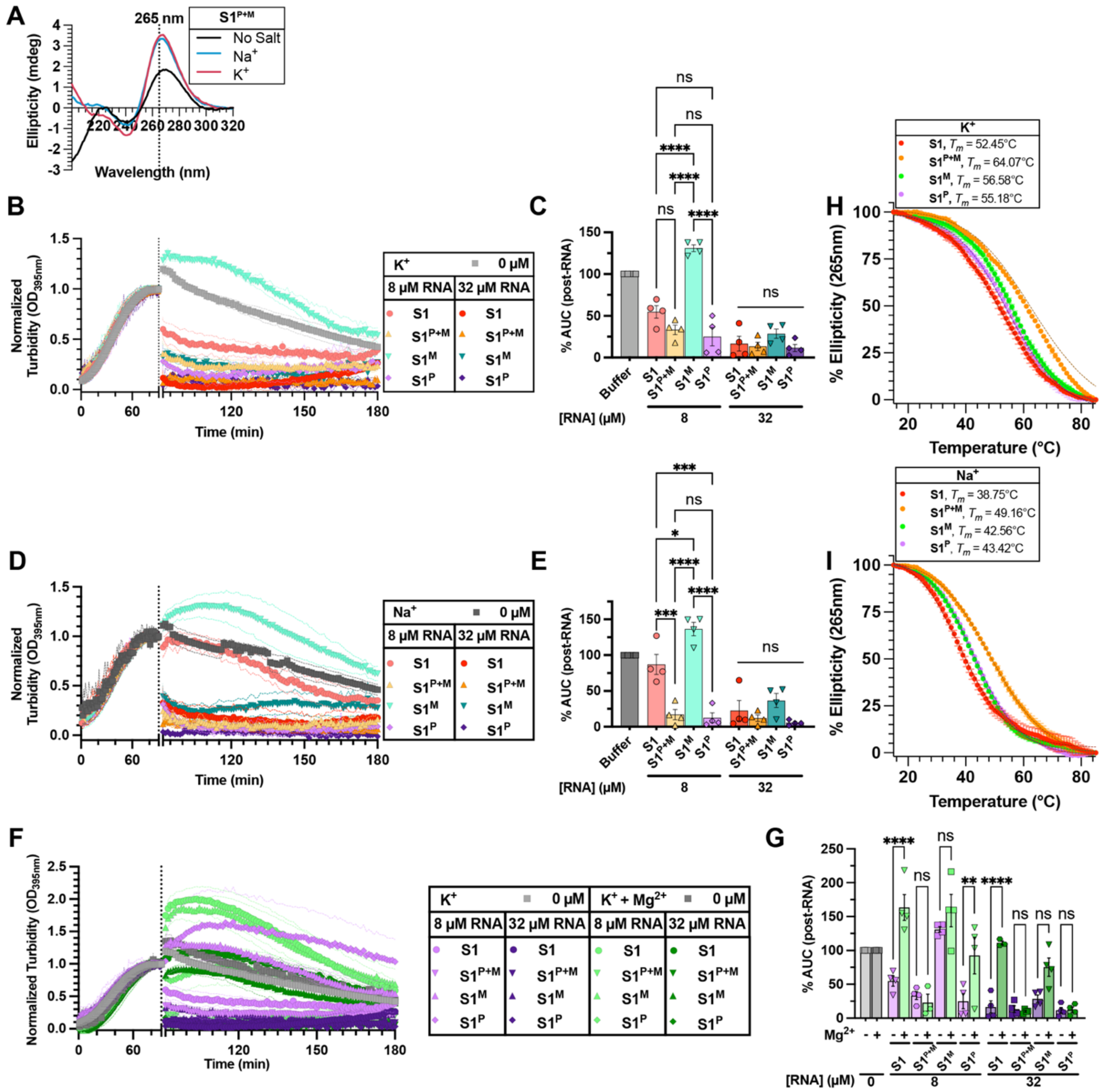

**Supplementary Figure 6. Chemical modifications enhancing rG4 stability strengthen RNA activity against FUS phase separation.**

(**A**) Representative CD spectra of RNA S1$^{P+M}$ collected at 25°C in indicated buffer.

(**B, D**) Turbidity measurement of FUS LLPS reversal with indicated RNA in TAB-K$^{+}$ (**B**) or TAB-Na$^{+}$ (**D**). LLPS of FUS (2 μM) was initiated, and monitored by turbidity at 395 nm. After 90 min, data collection was paused to add RNA. Data represent mean±SEM (n=4).

(**C, E**) AUC was calculated from turbidity curves in Fig. S6B (**C**) or Fig. S6D (**E**) after RNA was added. Data represent mean±SEM (n=4). Ordinary one-way ANOVA with Šidák's multiple comparisons test was used to compare the means within each concentration (ns P>0.05, *P≤0.05, ***P≤0.0005, ****P<0.0001).

(**F**) Turbidity measurement of FUS LLPS reversal with indicated RNA in TAB-K$^{+}$ ± Mg$^{2+}$. FUS (2 μM) LLPS was initiated, and monitored by turbidity at 395 nm. After 90 min, data collection was paused to add RNA diluted in 10 mM Tris-HCl pH 7.4 supplemented with 50 mM KCl (purple symbols) or 50 mM KCl + 3 mM $MgCl_2$ (green symbols). Data represent mean±SEM (n=3-4).

(**G**) AUC was calculated from turbidity curves in Fig. S6F after RNA was added. Data represent mean±SEM (n=3-4). Ordinary one-way ANOVA with Šidák's multiple comparisons test was used to compare the means (ns P>0.05, **P≤0.005, ****P<0.0001).

(**H, I**) Thermal melt curves of RNA S1 (red), $S1^{P+M}$ (orange), $S1^{M}$ (green), or $S1^{P}$ (purple) in $K^+$ (**H**) or $Na^+$ (**I**) collected at 265 nm. Data represent mean±SEM (n=3). Dashed lines represent the fitted curves used to obtain $T_m$.

Supplementary Figure 7

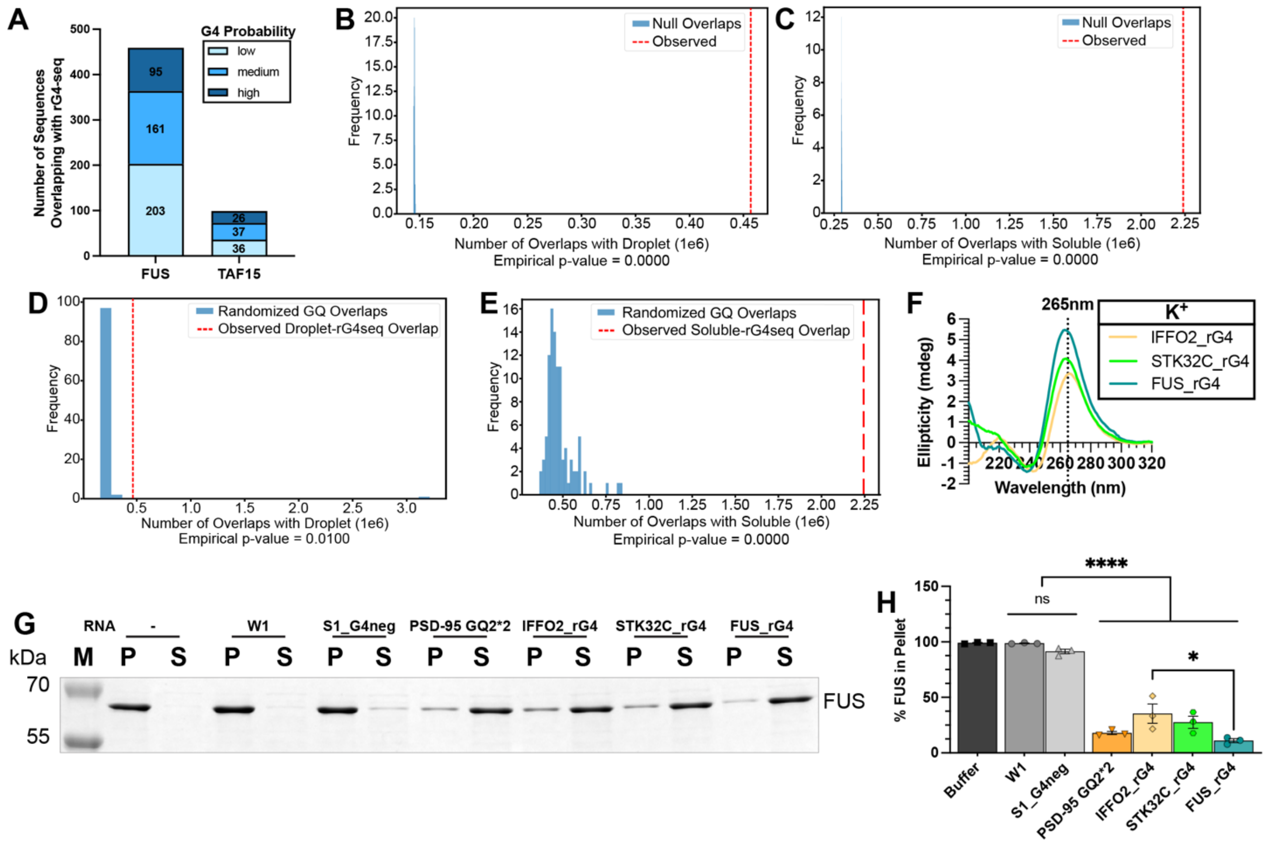

**Supplementary Figure 7. Transcriptome-wide bioinformatic analysis identifies novel RNA inhibitors of FUS LLPS and aggregation.**

(**A**) Number of sequences shared between FUS CLIP[3] or TAF15 CLIP[4] and rG4 database[5]. Overlap sequences with high probability of forming rG4 are in dark blue, those with medium probability are in blue, and those with low probability are in light blue.

(**B-C**) Permutation testing querying whether FUS droplet interactome (**B**) or FUS soluble interactome (**C**) specifically overlap with the rG4 database. rG4 regions were overlapped with randomly-generated regions from hg38 human transcriptome. Each randomly-generated file contained a number of regions consistent with the FUS droplet interactome or FUS soluble interactome files. This testing was repeated 100 times each for (**B**) and (**C**). Overlaps for each of the 100 tests were recorded (blue) and compared to the observed overlap between FUS droplet or soluble interactomes and rG4 database (red line).

(**D-E**) Permutation testing querying whether the rG4 database specifically overlaps with the FUS droplet interactome (**D**) or FUS soluble interactome (**E**) more than other regions. FUS droplet (**D**) or soluble (**E**) interactomes were tested against randomly-generated 30nt regions from the human transcriptome. These randomly-generated files were chromosome-matched and consistent with the number of regions originally in the rG4 database. Each test was repeated 100 times for FUS droplet and soluble interactomes. Overlaps for each of the 100 tests were recorded (blue) and compared to the observed overlap between FUS droplet or soluble interactomes and rG4 database (red line).

(**F**) Representative CD spectra of indicated RNAs collected at 25°C.

(**G**) Representative sedimentation analysis gels. FUS (5 μM) aggregation was initiated as in Fig. 6H. After 120 min, RNAs (100 ng/μL) were added and incubated at room temperature. Samples were collected 120 min after the addition of buffer or RNA and analyzed by sedimentation followed by SDS-PAGE.

(**H**) Quantification of SDS-PAGE images from Fig. S7G. FUS bands were quantified in pellet and supernatant fractions, and % FUS in pellet is reported. Data represent mean±SEM (n=3). Ordinary one-way ANOVA with Tukey's multiple comparisons test was used to compare the means between the conditions. (ns $P>0.05$, *$P\leq0.05$, ****$P<0.0001$).

#### SUPPLEMENTARY TABLES

**Supplementary Table 1. RNA oligonucleotides used in study.** QGRS Mapper[1] was used to predict whether RNAs would form rG4.

| Oligonucleotide | Sequence (5' → 3')<br><br>mX = 2'-O-methylation base<br>X(*) = phosphorothioate backbone | Length | Predicted to Form G4? | Oligonucleotide Reference |
|---|---|---|---|---|
| S1 | CUAGGAUGGAGGUGGGGAAUGG UAC | 25 | Yes | Guo et al.[6] |
| S1$^{P+M}$ | mC(*)mU(*)mA(*)mG(*)mG(*)mA(*) mU(*)mG(*)mG(*)mA(*)mG(*)mG(* )mU(*)mG(*)mG(*)mG(*)mG(*)mA( *)mA(*)mU(*)mG(*)mG(*)mU (*)mA(*)mC | 25 | Not Tested | This Paper |
| S1$^{M}$ | mCmUmAmGmGmAmUmGmGmAm GmGmUmGmGmGmGmAmAmUmG mGmU mAmC | 25 | Not Tested | Guo et al.[6] |
| S1$^{P}$ | C(*)U(*)A(*)G(*)G(*)A(*)U(*)G(*)G (*)A(*)G(*)G(*)U(*)G(*)G(*)G(*)G(* )A(*)A(*)U(*)G(*)G(*)U (*)A(*)C | 25 | Not Tested | This Paper |
| S1*2 | CUAGGAUGGAGGUGGGGAAUGG UACCUAGGAUGGAGGUGGGGAA UGGUAC | 50 | Yes | This Paper |
| S1*10 | GGCUAGGAUGGAGGUGGGGAAU GGUACCUAGGAUGGAGGUGGGG AAUGGUACCUAGGAUGGAGGUG GGGAAUGGUACCUAGGAUGGAG GUGGGGAAUGGUACCUAGGAUG GAGGUGGGGAAUGGUACCUAGG AUGGAGGUGGGGAAUGGUACCU AGGAUGGAGGUGGGGAAUGGUA CCUAGGAUGGAGGUGGGGAAUG GUACCUAGGAUGGAGGUGGGGA AUGGUACCUAGGAUGGAGGUGG GGAAUGGUAC | 252 | Yes | This Paper |
| S1_G4neg | GGCGUGAGAGUAGCGAUGUGUA GAG | 25 | No | This Paper |
| S2 | GAGGUGGCUAUGGAGGUGGCUA UGGAGGUGGCUAUGGAGGUGGC UAUG | 48 | Yes | Guo et al.[6] |
| S2_G4neg | GAGGUGCGUAGUGAGGUGCGUA GUGAGGUGCGUAGUGAGGUGCG UAUG | 48 | No | This Paper |
| W1 | UCAGAGACAUCAUCAGAGACAU CA | 24 | No | Guo et al.[6] |
| W1*2 | UCAGAGACAUCAUCAGAGACAU CAUCAGAGACAUCAUCAGAGAC AUCA | 48 | No | This Paper |
| TERRA | UUAGGGUUAGGGUUAGGGUUAG GG | 24 | Yes | Wang et al.[7] |
| $(GGU)_8$ | GGUGGUGGUGGUGGUGGUGGUG GU | 24 | Yes | This Paper |

| | | | | |
|---|---|---|---|---|
| $(G_4C_2)_4$ | GGGGCCGGGGCCGGGGCCGGGG CC | 24 | Yes | Fujino et al.[8] |
| PSD-95 | GGGGAAAAGGGAGGGAUGGG | 20 | Yes | Ishiguro et al.[9] |
| CaMKIIα | UGGGGGGGGCGGGUGGGAUGGG A | 23 | Yes | Ishiguro et al.[9] |
| Shank1a GQ | GGGGUUGGGGAGGGUGUAGGGG GUGGGG | 28 | Yes | Imperatore et al.[10] |
| PSD-95 GQ2*2 | GGGAGGGAGGGUGGGGGGAGGG AGGGUGGG | 30 | Yes | Imperatore et al.[10] *Doubled length of oligo |
| IFFO2_rG4 | GCAGGAAAUUCAGAAAUGUGGU GGGUGGGG | 30 | Yes | Kwok et al.[5] |
| STK32C_rG4 | GUCAAGGGCAUGGGUUGGGGUA GUGGGUGG | 30 | Yes | Kwok et al.[5] |
| FUS_rG4 | GGAGGCUUCCGAGGGGGCCGGG GUGGUGGG | 30 | Yes | Kwok et al.[5] |

**Supplementary Table 2. QGRS Mapper Results of previously-identified RNA inhibitors.** QGRS Mapper[1] was used to predict whether RNAs would form rG4. "S" denotes a strong inhibitor, "W" denotes a weak inhibitor, and "N" denotes no activity. If RNA is predicted to form rG4, major position(s), length(s), and G-Score(s) are reported.

| Oligonucleotide | Sequence (5' → 3') Predicted rG4 sequence is underlined | Predicted to form rG4? | rG4 Position | rG4 Length | rG4 G-Score |
|---|---|---|---|---|---|
| S1 | CUAGGAUGGAGGUGGGGAAUGGUAC | YES | 4 | 13 | 20 |
| S2 | GAGGUGGCUAUGGAGGUGGCUAUGGAGGUGGCUAUGGAGGUGGCUAUG | YES | 6 | 20 | 21 |
| | | | 27 | 14 | 18 |
| S3 | AUUGAGGAGCAGCAGAGAAGUUGGAGUGAAGGCAGAGAGGGGUUAAGG | YES | 23 | 26 | 21 |
| S4 | ACCAUGAUCACGAAGGUGGUUUUCCCAGGGCGAGGCUUA | YES | 15 | 21 | 14 |
| S5 | CUCCGGAUGUGCUGACCCCUGCGAUUUCCCCAAAUGUGGGAAA | NO | | | |
| S6 | AAAAAAAAAAAAAAAAAAAAAAAAAGUGAAGGCAGAGAGGGGUUAAGG | YES | 31 | 18 | 15 |
| S7 | GACUGAAAAAGGUGGGUUUCUUUU | NO | | | |
| S8 | AUUGAGGAGCAGCAGAGAAGUUGGAAAAAAAAAAAAAAAAAAAAAAAA | NO | | | |
| W1 | UCAGAGACAUCAUCAGAGACAUCA | NO | | | |
| W2 | GAAAAUUAAUGUGUGUGUGUGGAAAAUU | NO | | | |
| W3 | UUGUAUUUUGAGCUAGUUUGGUGAU | NO | | | |
| W4 | GGUGAGCACAGAGGUGAGCACAGA | NO | | | |
| W5 | CCAAUCUUCCUUCCAAUCUUCCUU | NO | | | |
| W6 | GAUGGAUUCCAGGAUGGAUUCCAG | NO | | | |
| W7 | AAACGGUCUGAUAAACGGUCUGAU | NO | | | |
| W8 | AAAGCGGCGAUGAAAGCGGCGAUG | NO | | | |
| W9 | UAUUGAUCCGGUUAUUGAUCCGGU | NO | | | |
| N1 | CGCUGGCAUCCACGCUGGCAUCCA | NO | | | |
| N2 | ACAGUCCCCCGGACAGUCCCCCGG | NO | | | |

| | | |
|---|---|---|
| N3 | ACCGGCGAACCGGCGAACCGG CGA | NO |
| N4 | CAGUAUUAUUUCAGUAUUAU UUU | NO |
| N5 | CCAUCCAGUCUACCAUCCAGU CUA | NO |
| N6 | CCAGUCUGGCCCCCAGUCUGG CCC | NO |
